# Learning from the unknown: exploring the range of bacterial functionality

**DOI:** 10.1101/2022.11.28.518265

**Authors:** Yannick Mahlich, Chengsheng Zhu, Henri Chung, Pavan K. Velaga, M. Clara De Paolis Kaluza, Predrag Radivojac, Iddo Friedberg, Yana Bromberg

## Abstract

Determining the repertoire of a microbe’s molecular functions is a central question in microbial biology. Modern techniques achieve this goal by comparing microbial genetic material against reference databases of functionally annotated genes/proteins or known taxonomic markers such as 16S rRNA. Here we describe a novel approach to exploring bacterial functional repertoires without reference databases. Our *Fusion* scheme establishes functional relationships between bacteria and assigns organisms to Fusion-taxa that differ from otherwise defined taxonomic clades. Three key findings of our work stand out. First, bacterial functional comparisons outperform marker genes in assigning taxonomic clades. Fusion profiles are also better for this task than other functional annotation schemes. Second, Fusion-taxa are robust to addition of novel organisms and are, arguably, able to capture the environment-driven bacterial diversity. Finally, our alignment-free nucleic acid-based Siamese Neural Network model, created using Fusion functions, enables finding shared functionality of very distant, possibly structurally different, microbial homologs. Our work can thus help annotate functional repertoires of bacterial organisms and further guide our understanding of microbial communities.

## INTRODUCTION

Exploring the molecular functional capabilities of microbes is key to understanding their lifestyles and contributions to the biogeosphere cycles that run our world(1-6). Microbial communities are often analyzed by taxonomically categorizing their members, defining their functional capabilities, and using this knowledge as a proxy for the community’s overall functional abilities(7-10). DNA-DNA hybridization (DDH) has long been accepted as the experimental gold standard for taxonomic classification of newly sequenced organisms and reclassification of existing ones (11,12); although note that experimental error in establishing DDH could negatively impact species delineation(13). For an easier comparison, DDH could be approximated using 16S rRNA similarity and bacterial morphology and physiology (14-16). However, more recent approaches analyze genome sequence properties, such as average nucleotide identity (ANI) and multilocus sequence similarity(17-21). These sequence-based methods are thus the *de facto* gold standard of organism classification, promising taxonomic precision and simpler and cheaper experimental use.

Notably, the above methods adopt a primarily phylogenetic view of bacterial relationships, assessing microorganisms’ likely evolutionary lineage based on genetic similarity. Horizontal gene transfer (HGT), i.e. the exchange of genetic material across taxonomic lineages, complicates this approach to bacterial classification(22-24). HGT is the primary way for evolutionarily distant organisms to acquire similar functional capabilities encoded by similar sequences(25-27). Conversely, evolutionarily close sequence-similar organisms can functionally diverge under environmental pressure. Given a shift towards analyzing the functional capabilities of microbes(8,28-30), i.e. “What are they doing?” instead of “Who are they?”, one might ask the question “Are these bacteria functionally related?” as opposed to “Are they evolutionary cousins?” The former question can be answered well, if incompletely, by phenetic approaches based on, for example, differentiation of cell wall composition, guanine-cytosine content, and the presence of lipids amongst others(31,32). We propose that genome-inferred bacterial functional annotations may further improve the resolution of these methods.

We previously developed Fusion, a method for evaluating microbial similarities based on shared functionality encoded in their genomes(28,33). This approach revealed relationships between organism groups that are overlooked when using taxonomic or DNA similarity alone. Here, in addition to updating our classification scheme for a faster and more precise way of dealing with a flood of microbial genomes, we made three key observations: (1) We validated our functional classification of proteins by evaluating the ability of Fusion functions to recall existing bacterial taxonomy. The expansion of the bacterial dataset in comparison to earlier work, from 1.3K to 8.9k organisms – a seven-fold growth, did not equivalently increase the proposed number of Fusion functions (25% increase, 335k vs 434k functions). We thus suggest that a large fraction of existing bacterial functions has already been captured. (2) Evaluating the capability of Fusion functions to recapitulate bacterial taxonomy we noted that bacterial functional relationships are often different from genome or evolutionary associations. While we have touched on this in previous work(28,33), here we demonstrate the stability of a function driven classification scheme and suggest it as a means for annotating functional abilities of newly sequenced organisms. (3) In an effort to improve protein functional annotations, we trained a Siamese Neural Network (SNN)(34) to label two gene sequences as encoding proteins of the same Fusion function. Our model captures functional similarity signals unavailable to the alignment-based or structural-comparison schemes, highlighting previously unexplored relationships in the sequence-structure-function (dis)continuum. We note that a recent study by Leman et al(35) has made similar observations across the three types of protein similarities. We further note that our approach could potentially be optimized to label functional profiles of microbial metagenomes directly from sequencing reads, i.e. without the need of assembly or metagenomic binning(36-39).

## MATERIALS AND METHODS

### Microbial proteomes

We retrieved a set of microbial proteomes from GenBank (40,41) (NCBI public ftp - ftp.ncbi.nlm.nih.gov/genomes/genbank/bacteria; February 28, 2018) and extracted the corresponding coding sequences from the complete bacterial genome assemblies. As per NCBI, complete assemblies are complete gapless genomic assemblies for all chromosomes, i.e. in bacteria, the circular genome and any plasmids that are present. Our resulting dataset thus contained the proteomes of 8,906 distinct bacterial genome assemblies with a total of 31,566,498 proteins (*full protein set*). We further redundancy reduced this set at 100% sequence identity over the complete length of the two proteins using CD-Hit (42,43). Our *sequence-unique protein set* contained 15,629,432 sequences. Sequences shorter than 23 amino acids (1,345 sequences) were removed from the set as this length is insufficient to determine functional similarity between proteins (44). All further processing was done on the resulting set of 15,628,087 sequences. Of these, 12.78M were truly unique, i.e. proteins for which no 100%-identical sequence exists in the original full protein set; the remaining 2.85M sequences represented the nearly 16M proteins that were redundant across organisms in our set.

### Computing protein functional similarities

Functional similarities between our sequence-unique proteins were assessed using HFSP(44). HFSP values reflect how likely two proteins perform the same function. Protein pairs scoring HFSP≥0 are assumed to be of the same function, with higher values indicating higher certainty of same function assignment (maximum HFSP =72). Specifically, we generated a set of all-to-all alignments with MMSeqs2(45) (evalue ≤ 1e-3, inclusion evalue ≤ 1e-10, iterations = 3). Note that due to the specifics of MMSeqs2, the two alignments for a every pair of proteins P_i_ and P_j,_ i.e. P_i_-to-P_j_ and P_j_-to-P_i_, are not guaranteed to be identical and thus may have different HFSP scores. We chose to conservatively represent each protein pair by only one, minimum, HFSP value. For every protein pair, we retained in our set only the alignments where this HFSP value was ≥0; at this threshold HFSP correctly predicts functional identity of proteins with 45% precision and 76% recall (44). Any protein without predicted functional similarity to any other protein in the sequence-unique protein set was designated as having a unique function, i.e. true singletons (766,050 proteins). Of these, 57,646 sequences represented 127,543 proteins in the full protein set, while 708,404 were truly unique. The remaining 14,862,037 proteins were connected by ∼22.2 billion functional similarities.

### Generating Fusion functions

We built a functional similarity network using the 22.2B similarities (edges) of the 14.86M proteins (vertices) as follows: For any protein pair P_i_P_j_, an edge was included if (1) HFSP(P_i_P_j_) was ≥ 30 or if (2) HFSP(P_i_P_j_) ≥ 0.7*max(HFSP(P_i_P_k_), HFSP(P_j_P_l_)), where proteins P_k_ and P_l_ are any other proteins in our set; note that P_k_ and P_l_ can but don’t have to be the same protein. The first cutoff at HFSP≥30, ensured that our protein pairs were often correctly assigned same function (precision = 95%). Our second criterion aimed to assuage the much lower recall (10%) and capture more distant relationships while introducing as little noise as possible, i.e. only reporting functionally similar pairs at specifically-targeted, stricter HFSP cutoffs. The resulting network contained 14,130,628 vertices connected by 780,255,934 edges (HFSP values used as weights); 731,409 proteins were disconnected from the network, i.e. *putative* functionally unique singletons. The network was composed of multiple connected components, where the largest contained 481,801 proteins (distribution of component sizes in Fig. S1).

We used HipMCL(46) (High-performance Markov Clustering), an optimized version of Markov Clustering(47,48), to further individually cluster the components of this network into functional groups. Note that as HipMCL requires a directed graph as input, we converted each edge in our data into a pair of directed edges of the same weight. The key parameters chosen for each HipMCL run were S=4000, R=5000, and inflation (I) =1.1. This clustering resulted in 1,432,643 protein clusters as well as 1,235 clusters containing only one protein, i.e. additional *putative singletons* for a total of 732,644.

Given the high functional similarity threshold used above, these clusters were exceedingly specific. We aimed to generate somewhat broader functional definitions to avoid artificially inflating functional diversity. We thus extracted representative proteins of each cluster that were further re-clustered using lower functional similarity thresholds. Representative sequences of each of the 1,432,643 MCL clusters were extracted using CD-Hit at 40% sequence identity (with default parameters and selecting the longest protein per CD-HIT cluster). Note that only 7% of the MCL clusters had multiple representative sequences; thus, a total of 1,632,986 cluster representatives were collected. To this set of representatives, we added the putative singletons for a total of 2,365,630 proteins. These were used to generate a new functional similarity network by including all edges with HFSP(P_i_P_j_)≥0. Note that 226,346 (∼10%) of these were not similar to any other representative proteins; of these, ∼40k were originally designated putative singletons. The resulting functional similarity network comprised 2,139,284 vertices and ∼303M edges. The network was re-clustered with HipMCL (S=1500, R=2000, I=1.4; smaller inflation values did not generate results due to MPI segmentation faults that could not be resolved) generating 433,891 Fusion functions.

### Enzymatic function annotation

We evaluated shared enzymatic functionality of proteins using Enzyme Commission (EC) annotations (49). The EC Number is a numerical classification scheme for enzymes, based on the chemical reactions they catalyze. Every EC consists of four numbers (separated by a period), indicating the corresponding class and sub-classes of the enzyme. Enzymes annotated with the same EC number at all four levels are considered functionally identical, although EC identity at three highest levels may be considered sufficient for informing functional similarity.

Information about protein enzymatic activity was extracted from Swiss-Prot(50,51) (June 2021) as follows: for each protein there had to be (1) experimental evidence for protein existence at protein level, (2) experiment-based functional annotation, and (3) only one EC number, fully resolved to all four levels. The resulting dataset was redundancy reduced at 100% sequence identity across the entire protein length. Swiss-Prot entries sharing the same sequence, but assigned different EC annotations, were excluded from consideration. The final data set contained 18,656 unique proteins and 4,269 unique EC annotations. The overlap between the EC data and the Fusion protein set (*Fusion enzyme set*) comprised 4,206 unique proteins in 1,872 unique EC annotations.

### Pfam data

Protein mappings to Pfam(52) domains (Pfam-A version 34) were generated using pfamscan v1.4(53) with default values; in hmmscan(54) (hmmer v3.3), HMM evalue (-E = 10) and domain evalue (--domE = 10) were used. If the sequence hit multiple Pfam domains belonging to the same clan/family, only the clan was reported. For 12,720,756 sequence-unique proteins (85% of our 14.86M) the set of non-overlapping Pfam domains and their order in sequence were extracted, e.g. given domains X and Y, the domain arrangements ‘XYY’, ‘XY’ and ‘YX’ are regarded as three individual occurrences; the remaining 15% of the proteins did not match any Pfam-A domain. We thus identified 92,321 unique Pfam domain arrangements. These corresponded to 58,021 domain sets, where the domain arrangements ‘XYY’, ‘XY’ and ‘YX’ resolve to only one domain set representation (X,Y).

### Overlap between Fusion clusters and GTDB

In order to compare Fusion functions to the set of 120 marker proteins/protein families that GTDB uses (TIGRFAM & Pfam families) to establish taxonomic relationships between organisms (bac120), Fusion proteins were associated with TIGRFAM (release 15.0 – September 2014) & Pfam (PFAM-A version 34) domains using hmmscan (hmmer v3.3) at default thresholds (hits with HMM evalue, -E = 1 and domain evalue, --domE = 10). Only one best TIGRFAM/Pfam hit (i.e. smallest e-value) was extracted per protein. Fusion functions were assigned the set of TIGRFAMs/Pfams according to their proteins matches. Finally, the overlap between domain associations of Fusion functions and the TIGRFAMs/Pfams used by GTDB as marker genes was evaluated.

### GeneOntology annotations

GO(55,56) “molecular function” annotations were extracted from the GO 2021-09-01 release. For each protein, its set of GO annotations included all protein self-annotations, as well as annotations of its parent nodes, i.e. other nodes connected via an “is a” edge up to the root of the molecular function subgraph. This resulted in 25,825 sets of GO terms for 7,313,428 (49% of 14.9M) sequences-unique proteins.

### Comparing Fusion functions to existing functional annotations

We compared Fusion functions to EC and Pfam annotations by calculating the homogeneity (h, Supplementary Eqn. S1), completeness (c, Supplementary Eqn. S2) and V-Measure (v, Supplementary Eqn. S3) (57) values using scikit/python (58). When comparing Fusion functions to, for example, EC numbers, homogeneity describes how often a Fusion function is associated with multiple EC numbers. That is, a high homogeneity (close to 1) signifies a clustering where most Fusion functions have an association to only one EC number. Completeness describes how often a specific EC number can be found in different Fusion functions. A high completeness (close to 1) indicates that for most ECs, a specific EC number is associated with only one or a small number of functions. V-Measure represents the harmonic mean between homogeneity and completeness. A V-measure of 1 is indicative of an optimal clustering, where each function is only associated with one EC number, and an EC number is only associated with this one function.

### Taxonomy information

Our taxonomic analyses were conducted on the basis of two taxonomy schemes: the NCBI taxonomy(59) and the GTDB(60) (genome taxonomy database). NCBI taxonomy rank information for each assembly was retrieved during protein dataset extraction (Feb 2018) and is available for all 8,906 organisms in our set. GTDB taxonomy information was extracted from GTDB release rs202 (April 2021). Genbank assembly ids were mapped to bacterial assemblies available in GTDB. GTDB taxonomy information is available for 99% (8,817) of the organisms.

### Balancing the assembly set

According to GTDB, the 8,906 assemblies/organisms in our set belong to 3,005 species. Of these species, 2,252 (75%) have only one associated organism, whereas others have hundreds; e.g. *E. coli* and *B. pertussis* have 472 and 360 assemblies, respectively. We generated a balanced organism set to reduce this unevenness. First, we reduced our full set of 8,906 assemblies to retain the 3,012 genomes that were representative of strains included in the GTDB bac120 phylogenetic tree. Note that of these, 2,206 genomes were in both GTDB and our data, while 806 genomes were not present in our set and were represented by other assemblies of these same strains. Using dendropy(61), we then extracted from the full GTDB bac120 tree (47,895 organisms) a subtree containing only these 3,012 representatives while retaining the original branch lengths. We used Treemmer(62) to determine which leaves to retain in our set such that the RTL (relative tree length) of the pruned tree was ≥0.90. RTL is used as an indicator of retained genetic diversity after pruning, reflected as the sum of all branch lengths in the pruned tree in relation to the full tree. We thus selected 1,502 assemblies (further referenced to as the *balanced organism set*) – a minimum set of organisms that retains at least 90% genetic diversity present in our complete set of 8,906 assemblies.

### Computing organism functional similarity

Each organism in our set can be represented by a functional profile, i.e. a set of corresponding Fusion functions, Pfam domains, or GO annotations. Functional similarity between the functional profiles of two organisms, F_i_ and F_j_, was calculated, as previously described (28,33), by dividing the number of their shared functions by the size of the larger of the two profiles (Eqn. 1).

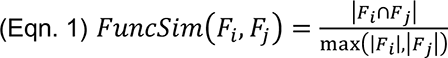

Fusion functional profiles for similarity calculations were generated at Fusion Level 1 with and, separately, without the inclusion of singletons. Pfam functional profiles were generated using Pfam domain arrangements and, separately, domain sets, as described above. GO functional profiles were generated using the GO terms extracted per proteins as described above. Note that Pfam and GO annotations are not available for all proteins, but each protein has an associated Fusion function. Thus, each method-based functional profiles (i.e. GO vs Pfam vs Fusion) of a single organism could be based on different sets of proteins.

We computed the precision/recall (Eqn. 2) values for correctly identifying two organisms as being of the same taxonomic rank based on their shared functional similarity. This was done at each taxonomic rank (phylum, class, order, family, genus, species) for both taxonomic definitions (NCBI and GTDB) and using a series of similarity thresholds ranging from 0 to 1 in increments of 0.01.

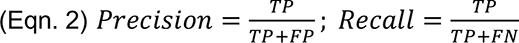

Here any pair of two organisms of the same taxonomic classification above the chosen threshold are true positives (TP), whereas pairs below the threshold are false negatives (FN). Any pair of two organisms of different taxonomic classifications above the similarity threshold are false positives (FP), while pairs below are true negatives (TN).

### Grouping organisms by functional similarity

An organism similarity network was generated using Fusion functional profiles. Here assemblies (vertices) were connected by Fusion functional similarity edges; the resulting network is complete (all-to-all edges are present) as any two organisms share some similarity. We used Louvain clustering (63) to identify organism groups; implemented in ‘python-louvain’ (https://github.com/taynaud/python-louvain), an extension to ‘networkx’ (https://networkx.org). Organism groups at varying levels of granularity were generated by varying the resolution threshold parameter of Louvain clustering (resolution 0 to 1.5 in increments of 0.01), where larger resolution values lead to fewer but larger clusters. The V-measures (Eqn. 3) of the resulting partitions (“predicted labels") vs. GTDB taxa (reference labels) were calculated.

### 16S rRNA extraction and similarity calculations

16S rRNA sequences were extracted from the NCBI GenBank database for 8,479 of the 8,906 organisms (427 organisms were missing annotated 16S rRNAs). From RDP (Ribosomal Database Project, v11.5)(64), we further extracted all 16S rRNA sequences and their corresponding multiple sequence alignment (MSA). The 16S rRNAs of the 8,479 organisms that were not contained in the RDP MSA were added using Infernal 1.1.4 (65) and the RDP bacterial covariance model. Using the resulting MSA we extracted gapless pairwise sequence identities for all 16S rRNA pairs (i.e. 683,261,061 pairs between 36,967 16S rRNA sequences).

We calculated the optimal F-measure (Eqn. 3) for both identifying organisms of the same species/genus using measures of 16S rRNA identity and Fusion organism similarity (Eqn. 1). Here, true positives (TP) are organisms of same taxon, attaining an identity or similarity measure at or above the chosen threshold, false negatives (FN) are organisms of same taxon but scoring below the threshold, and false positives (FP) are organisms of different taxa and scoring at or above the threshold.

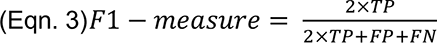

### MASH organism similarity calculations

For each organism in our dataset we extracted pre-computed k-mer hash sketches using the MASH(66) -provided RefSeq genome sketch database containing 126,381 NCBI prokaryotic genomes. Jaccard Distances between organism sketches were calculated according to Ondov et.al. Similar to the 16S analysis, we calculated the optimal F-measure (Eqn. 3), Precision, and Recall (Eqn. 2) for identifying two organisms of same species or genus.

### Machine learning-based predictor of shared protein functionality

We trained a Siamese Neural Network (SNN) (34) predictor to assess whether any two DNA sequences encoded proteins of the same Fusion function. SNNs are a class of neural network architectures that contain two identical subnetworks, i.e. the networks have the same configuration with the same parameters and weights. This type of network is often used to find the similarity of the inputs – in our case, two sequences encoding proteins of the same function. Because SNNs identify similarity levels, rather than predicting specific classes of each input, they require significantly less data for training and are less sensitive to class imbalance. The latter was particularly a benefit here because the number of sequence pairs of different functions necessarily drastically exceeds the number of pairs of the same function. Additionally, as SNNs output a similarity metric rather than a probability score, they are likely specifically informative of the various levels of functional similarity, e.g. for a given pair of enzymes, whether two genes act upon the same bond vs. whether they use the same electron donor.

To train the model, we extracted 70 random Fusion functions, each containing at least ten different proteins from our sequence-unique set. The set of functions was split 50/10/10 for training, testing and validation. For training and validation, we balanced the dataset, i.e., we randomly selected gene sequence pairs such that 50% of the pairs included genes of same Fusion function and 50% were of different function. The final training set contained 20M gene sequence pairs generated from 29,907 sequences, the validation set contained 200,000 pairs and 9,982 sequences respectively. In testing we used balanced as well as imbalanced data sets. The imbalanced test set was generated to better resemble real-world data with a split of 90%/10% where 90% of the sequence pairs are between sequences of different function. The test set contained 100,000 sequence pairs generated from 1,000 gene sequences.

We tokenized protein-encoding genes to codons, i.e. split into non-overlapping 3-nucleotide chunks of sequence and projected each token into the LookingGlass(67) embedding space (length=104). The embeddings were then processed via an LSTM (68) and further used in SNN training. Note that at most the first 1,500 tokens were embedded per sequence. For sequences shorter 1,500 codons, the embedding vector was zero padded, i.e., any position in the vector after the last token was set to 0. The model was trained and validated in 50 iterations on our balanced training/validation data set. After 50 iterations performance of the model reached a precision of 0.72 and recall of 0.72 on the validation set at the default threshold of 0.5. The final model was tested on the imbalanced (90/10 split different/same function sequence pairs) attaining a precision of 0.22 and recall of 0.80 at the default prediction score cut-off of 0.5.

To further evaluate the model, we extracted a set of Fusion functions associated with only one level 4 EC annotation, but where the EC annotation was associated with multiple Fusion functions. We then predicted SNN scores for three sets of protein pairs: (1) proteins from the same Fusion function and same EC annotation, (2) proteins from different Fusion functions and same EC annotation, and (3) proteins of different Fusion functions and different EC annotation.

### Structural alignments of Fusion proteins

We extracted from the PDB(69,70) (May 2022) the available structure information for proteins in our set, i.e. 79,464 chains/entities mapping to 5,153 protein sequences in our sequence-unique protein set. Where multiple PDB structures mapped to one protein sequence we selected the PDB entry with the best resolution (lowest Å). For this set, we used foldseek(71) (–alignment-type 1, --tmscore-threshold 0.0) to identify structure pair TMscores(72) from TM-align(73). When a protein sequence pair resolved to multiple PDB entity (chain) pairs we selected the entity pair with the highest TMscore. Note that Foldseek was unable to generate TMscores for 498 PDB structures (mapping to 1,005 protein sequences) due to computational limitations and we excluded any structural/protein pair that included one of these from further consideration.

For the resulting 8,600,878 protein pairs we generated SNN prediction scores. For 8,150,441 of 8,600,878 (95%) pairs no TM-scores could be generated as they did not pass the pre-filtering step of Foldseek, i.e. they had no similar folds at all; for these we assumed a TMscore = 0. Notably, 144,829 (1.7%) of these were still predicted by the SNN to have high functional similarity (SNNscore≥0.98); we assume this percentage to be the approximate error rate of the SNN.

We also created subsets of PDB entity pairs where each protein was annotated with an E.C. number, i.e. proteins extracted for the Fusion enzyme set.

## RESULTS

### Fusion reflects and augments known functionality

After removing identical sequences (Methods, Supplementary Information), we computed (using HFSP (44), Homology-derived Functional Similarity of Proteins) functional pairwise similarities (edges) between proteins (vertices) and clustered the resulting protein similarity network to determine the molecular functions likely carried out by proteins in our set (Methods; Fig. 1). Note that in our earlier work (28,33) to establish microbial functionality we used different edge-weighing and clustering techniques over a significantly smaller number of proteins (4.2M earlier vs. 31.5M in this work) and organisms (1,374 vs. 8,906). Our earlier approach yielded ∼900K singletons and ∼335K functional groups of more than one protein. Here, we significantly expanded our ability to capture protein functionality, obtaining only a few more singletons (∼947K) and 25% more (433,891) clusters of functionally similar proteins, dubbed *Fusion functions*, ranging in size from two to 118,984 proteins (Fig. S1). The limited difference in numbers of identified functions suggests that we have already explored much of the possible bacterial functional space.

**Fig. 1.**
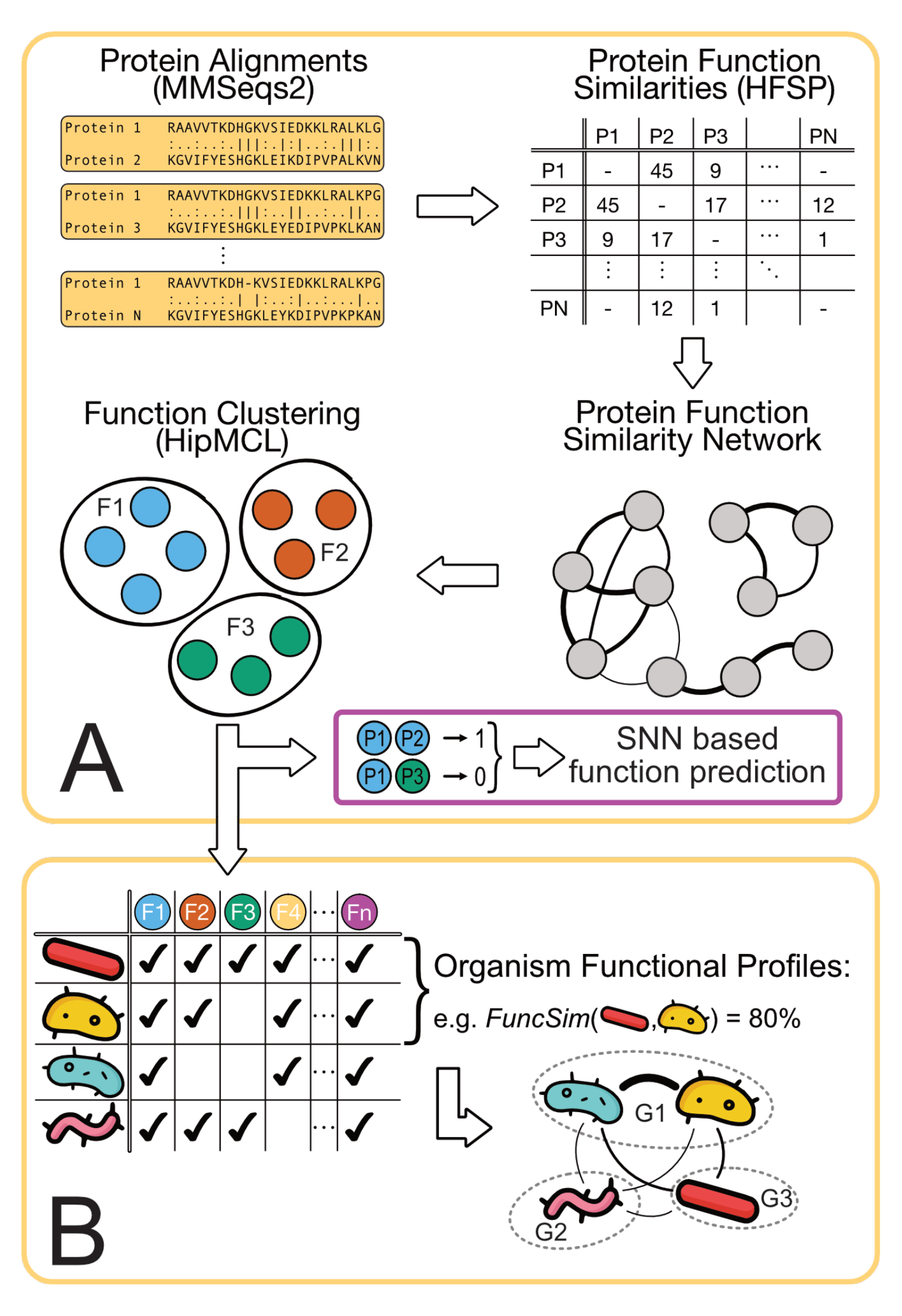
Fusion workflow. **(A)** Fusion functions are formed by generating all-against-all MMSeqs2 protein alignments between all ∼15.6 million proteins in our dataset and establishing protein functional similarities from these alignments using HFSP. The resulting protein function similarity network, where proteins are nodes and the similarity is the weight of the edge or no edge if no similarity was found, is clustered with HipMCL and function clusters are retrieved. The pairs of proteins within the same cluster were used as input for SNN training. **(B)** Furthermore, vectors of functions (rows) represented organisms in our set. Organism Functional Profile Similarities were computed as row-wise vector comparisons. These were used to generate an organism similarity network, where nodes are organisms and edges are weighed according to functional similarity. Clustering this network at different thresholds yielded the Fusion functional taxonomy.

This collection of Fusion functions, particularly the large number of functions containing few proteins, is contrary to expectations of functional diversity as compared to, e.g. 19,179 Pfam-A families/clans (Pfam v34, Methods) (52) and 11,185 molecular function GO terms (GeneOntology version 2021-09- 01; Methods) (55,56). This discrepancy between the annotations is likely due to definition of function.

Pfam-A, for example, needs many sequences per family to build multiple sequence alignments (MSAs) for Hidden Markov Model (HMM) construction; thus, some of our functions may simply have not contained enough sequences to recapitulate a Pfam family. Of the Fusion functions, only 15% (65,663) have at least 20 sequence-unique proteins, i.e. the lower limit for even the less-precise MSAs (74). Furthermore, Pfam domains are not functionally precise as the same domain is often reused in different functions (75-78) and one protein can have more than one domain; thus, one family is likely have more than one function. In fact, of these 66K functions (≥20 proteins per function), 80% (52,678) contain proteins with one or more non-overlapping Pfam domains (∼1.6 domains per protein; 10,114 unique domains total) and ∼11 Fusion functions per domain. Of the ∼370K smaller functions (<20 proteins), 128,128 have at least one Pfam-A domain. We hypothesize that the remaining ∼240K functions that are not identifiable by Pfam may be responsible for highly specific bacterial activity.

We calculated homogeneity (Supplementary Eqn. S1) and completeness (Supplementary Eqn. S2) for how well the Fusion functions (180,806 functions of >1 sequence) of the 12,611,237 proteins with at least one Pfam domain compared to Pfam-A domain assignments (Methods). We used measures of homogeneity and completeness; an optimal homogeneity (=1) would indicate that each function only contains proteins with one domain and an optimal completeness (=1) would indicate that all proteins with a specific Pfam domain only fall into a single function. Due to the absence of the one-to-one mapping between families and functions (discussed above), neither optimal completeness nor heterogeneity are possible for our data. However, both were high (homogeneity =0.9, completeness =0.79); that is, Fusion captured much of the Pfam-like functional diversity.

We further compared the Fusion functions with their respective Pfam domain sets, i.e., 57,165 collections of Pfam domains per protein without accounting for domain order in sequence. This comparison marginally increased completeness (=0.8) and homogeneity(=0.94), the latter suggesting that each Fusion function most often only contained proteins of one domain set. Additionally considering domain order (91,113 arrangements), we observed similar homogeneity (=0.93) and completeness (=0.81). The discrepancy between homogeneity and completeness indicates that while each Fusion function is highly specific to a given Pfam domain set or arrangements, each domain set/arrangement might encode multiple functions.

The above finding is in line with the observation that Pfam domain arrangements do not always report experimentally defined functionality(79). Here the precision of Fusion functions is important. For example, the *Geobacter sulfurreducens* acyltransferases (R)-citramalate synthase (AAR35175, EC 2.3.1.13) and *Salmonella heidelberg* 2-isopropylmalate synthase (ACF66296, EC 2.3.3.182/2.3.3.21) have the same domain arrangement (HMGL-like pyruvate carboxylase domain, PF00682, followed by a LeuA allosteric dimerization domain, PF08502) but have a different 4^th^ digit Enzyme Commission classification (EC) number (49), indicating their different substrate specificities. Notably, these proteins fall into two different Fusion functions.

To evaluate Fusion functional mappings more broadly, we collected the available experimentally derived EC annotations for proteins in our set (4,206 proteins, 1,872 unique EC numbers) and measured the similarity of these with the corresponding 1,893 Fusion functions. Fusion functions more closely resembled annotations of enzymatic activity (homogeneity = 0.95, completeness = 0.94) than those of Pfam domains. This finding suggests that our Fusion functions capture aspects of molecular function better than domain-based annotations.

In evaluating Fusion-based functional annotations, we were consistently faced with the inherently limited knowledge of microbial functionality. How can one evaluate the precision of Fusion function definitions at scale if no annotations of the component proteins exist? To address this question, we explored whether bacterial taxonomic classification, driven by genomic signatures as well as morphology and physiology, may be sufficiently informative of the organism functional repertoires. We suggest that functional annotations that reflect existing taxonomies may be deemed validated.

### Organism functional profiles capture taxonomy

For each organism of the balanced organism set, we extracted Fusion, Pfam-A domain arrangement, and GO term functional profiles. A functional profile is the set of functions of a single organism, e.g. the set of Pfam-A domain arrangements encoded by the proteins of that organism (Methods). On average, per organism Fusion, Pfam-A and GO term profiles were of size 2,133, 1,479, and 776, respectively (Fig. S2). For each organism pair, we computed profile similarity, i.e. the count of functions found in both profiles divided by the larger functional profile (Methods; Eqn. 1). On average, the (larger) Fusion-based functional profiles were less similar than the (smaller) Pfam and GO -based profiles (Fig. S3). A pair of organisms were predicted to be of the same or different taxon based on whether their similarity exceeded a set threshold ([0,1] in steps of 0.01). Predictions were compared against NCBI(59) and GTDB(60) taxonomies at six levels (phylum through genus; Methods). Note that we could not assess the species level, since no two organisms of the same species were retained in the balanced organism set.

As expected, all functional profiles were better than random at annotating microbial taxonomy (Fig. S4). Both Fusion and Pfam outperformed GO annotations in this task. Fusion profiles were better than Pfam, e.g. at 50% recall (Eqn. 2) of identifying two organisms of the same GTDB phylum, Fusion and Pfam achieved 75% and 48% precision (Eqn. 2), respectively. This advantage was also present across deeper taxonomic ranks. We note that Fusion’s improvement over Pfam did not stem from the difference in the number of functions per organism (profile size) as the predictive power of the functional profile size was only marginally better than random (Fig. S4). These findings confirm that organism similarity established via comparison of functional profiles carries taxonomy-relevant information. We thus further asked whether comparing functional capabilities can reveal organism relationships that phylogeny-based microbial taxonomy, muddled by horizontal gene transfer, is unable to resolve.

### Functional profiles are more informative of taxon identity than 16S rRNA

The genetic marker most frequently used for organism taxonomic classification is the 16S rRNA gene(15) – a non-coding gene that, by definition, cannot be captured by Fusion. To evaluate its predictive power, we extracted 16S rRNA sequences for each genome in our complete set and calculated sequence identity for all 16S rRNA pairs (Methods).

Sequence similarity between 16S rRNA pairs below 97% is generally accepted as an indication that the organisms are of different species(80). Indeed, we found that 98.7% (663.7M of 683.3M) of the 16S rRNA pairs that originate from different species (complete NCBI bacterial genomes collection; Methods) fall below the 97% sequence identity threshold, while only 2% of same species pairs do (Fig. 2, Fig. S5). That is, below this sequence identity threshold nearly all (99.96%) sequence pairs were of organisms of different species, confirming the 97% threshold as an excellent measure of organism taxonomic difference.

**Fig. 2.**
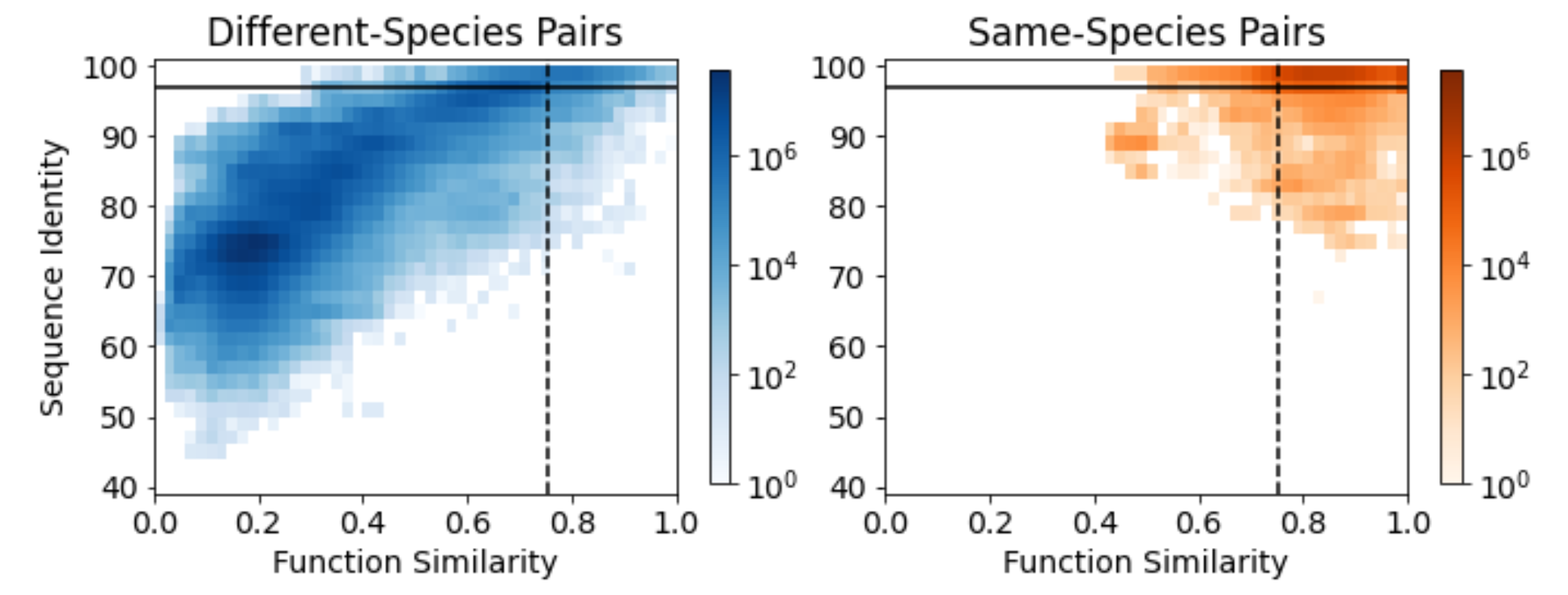
16S rRNA identity and functional similarity capture different taxonomic patterns. Density plots capture the location of pairs of different species (left, blue) and same species (right, orange) organisms in the space defined by the 16S rRNA identity (y-axis) and Fusion similarity (x-axis). Horizontal solid and vertical dashed lines represent the 16S rRNA and Function similarity thresholds of 97% and 75.5%, respectively.

Using the 97% sequence identity threshold as an indicator of taxon identity, however, is impossible. Many genomes have multiple 16S rRNA genes (81). In our set, 625 pairs of 16S rRNAs extracted from the same genome were less than 97% identical (minimum similarity =75.8%); in these cases, the marker gene similarity could not even identify the same genome, let alone same species. Furthermore, while almost all of same-species 16S rRNA pairs were ≥97% identical, nearly half of all pairs above this threshold belonged to different species (recall=98%, precision=55%, Fig. S6).

In contrast, at the optimal Fusion organism functional profile similarity threshold of 75.5% (Eqn. 1; threshold established via peak F1-measure, Eqn. 3; Fig. S7), organisms were correctly identified to be of the same species with 80% precision (recall=94%, Fig. S4). At a matched level of recall, function comparisons were also more precise than 16S rRNA (75% vs. 55% precision, at 98% recall). Furthermore, Fusion achieved 95% precision for more than a third (35%) of the organism pairs, whereas 16S rRNA measures were this precise for less than a fifth (17%). The ability of 16S rRNA to identify organisms of the same genus at the commonly used threshold of 95% also left much to be desired (43% precision, 78% recall). Fusion performance was significantly better (90% precision, 70% recall) when using optimal functional similarity threshold (72.3%) established for this task.

Functional profiles augmented 16S rRNA in determining organism species. For example, for all organism pairs sharing ≥97% 16S rRNA identity, additionally requiring a Fusion functional similarity of 75.5% lead to an increased precision of 86% vs. 55% for 16S rRNA or vs. 80% for Fusion similarity alone; recall was slightly decreased to 92% vs. 98% for 16S rRNA and 94% for Fusion alone. These findings suggest that functional similarity is somewhat orthogonal to 16S rRNA similarity in defining taxonomic identity and reaffirm the likely impact of Horizontal Gene Transfer on taxonomic classification.

We also note that the lack of precision in 16S rRNA-based comparisons has negative implications for metagenomic analysis, where 16S rRNA abundance is often used to assess sample taxonomic composition and functional diversity. Fusion functional annotations, on the other hand, can be optimized to target shorter sequencing reads e.g. via faser analysis(82), and thus infer a microbiome functional, if not taxonomic, composition directly from metagenomic sequencing experiments.

### Few functions are sufficient to accurately identify taxonomy

Fusion’s success in taxonomic classification is not unexpected. Using methods that take advantage of whole genome sequences, however parsed, is nearly guaranteed to be advantageous in comparison to 16S rRNA-based taxonomy determination(66,83,84). Exact comparison of genomes, however, is time-consuming. More useful approximations include methods like MASH(66) and Dashing(85), i.e. tools that use representative genome sequence sketches, and FastANI(84), which computes average nucleotide identities (ANI) across orthologous genes. We note that these methods are comparable in taxonomic classification accuracy, with ANI shining for species/strain differentiation, while sketch-based methods performing better at higher taxonomic levels(83,84). For our full set of organisms MASH achieved an optimal F1-score for same species identification at 96% k-mer sketch similarity (Figure S8; Methods). For the balanced organism set, MASH performed similarly to Fusion in terms of Precision/Recall at all taxonomic levels except genus (Figure S9). This result is in line with expectations given that whole genome comparisons should also capture functional similarity inferred from sequence comparison. However, sketch-based and ANI analyses lack the deeper understanding of the shared (or differing) bacterial functionality. They also do not lend themselves easily to the selection of taxonomic markers.

Earlier studies argue that a small number of carefully chosen marker genes/protein families are sufficient to determine taxonomic relationships of bacteria (60,86). However, to be comparable across organisms, these genes should be ubiquitously present. We investigated whether a subset of Fusion functions could correctly identify two organisms of the same taxon. To this end, we progressively subset the number of Fusion functions used to generate organism functional similarities (100k, 50k, 25k, 10k, 5k, and 1kfunctions). We used two approaches for function selection: (1) we chose the functions based on how frequently they appeared in the balanced organism set and (2) randomly sampled from the whole pool of functions. Importantly, our approach was based on the presence or absence of specific functional abilities encoded by these genes rather than their sequence similarity. We found that just 1,000 common (largest) Fusion functions were sufficient to classify organism pairs into the same taxon, competing with a “complete Pfam”-based approach (Fig. 3). The same was true for taxonomic levels of order through genus with a set of 5,000 randomly selected functions (Fig. S10).

**Fig. 3.**
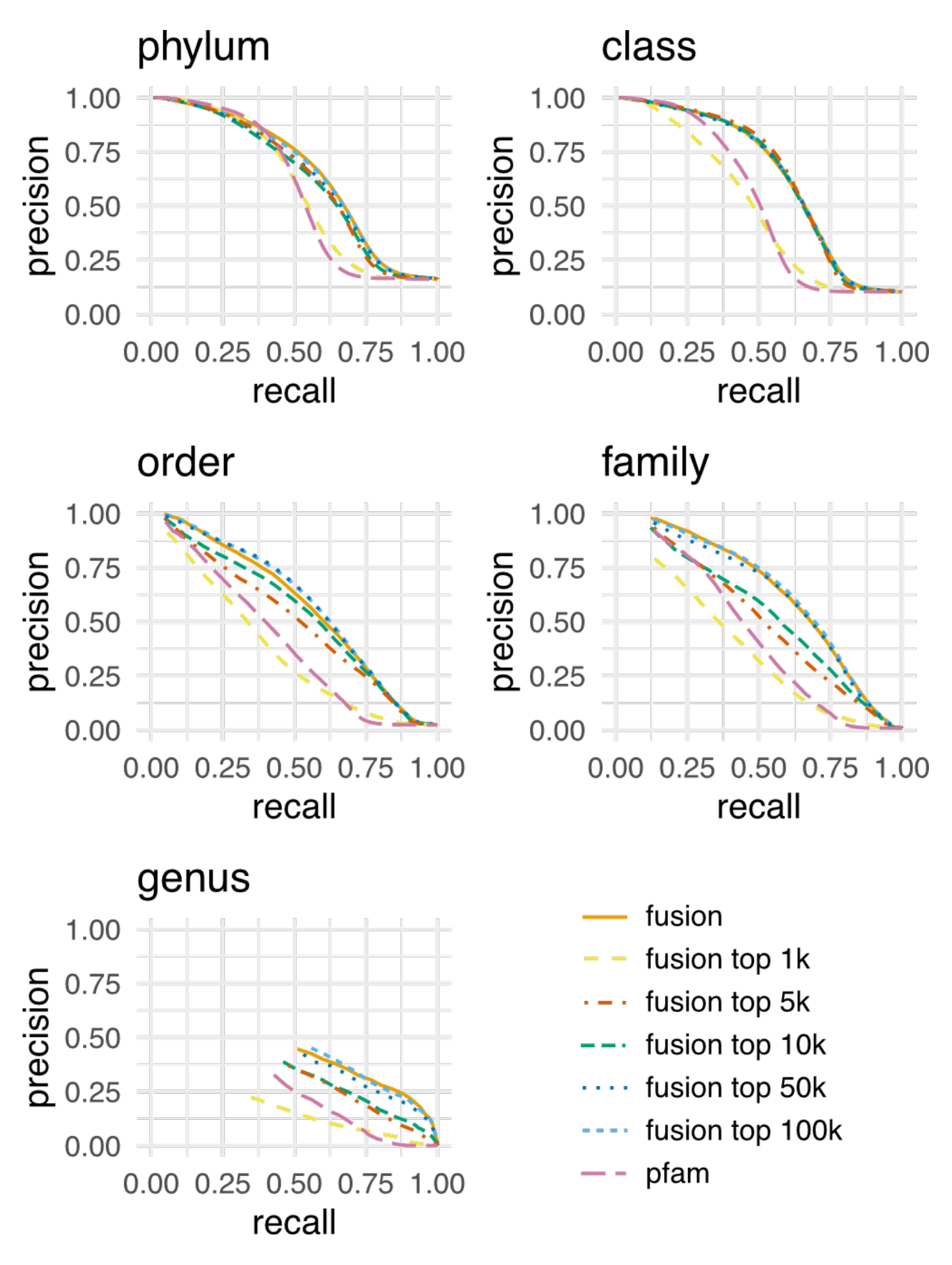
A subset of common fusion functions is sufficient to determine taxonomic relationships. Each panel describes the precision (y-axis) at a given recall (x-axis) for correctly identifying two organisms sharing the same taxonomic rank. Only precision/recall pairs where predicted positives pairs (TP & FP) make up at least 0.1% of all possible pairs are displayed. Sets of only the largest 100k (light blue) and 50k (dark blue) functions perform almost identically to all Fusion functions (solid red line) in distinguishing organism taxonomy at all taxonomic levels. For the largest 10k (olive) and 5k (red) functions, the same holds true for phylum and class, whereas order through genus performance is slightly reduced. Notably the largest 1,000 functions (yellow) perform similar to a complete Pfam domain arrangement-based approach (purple).

We further evaluated the overlap between the selected Fusion functions and the marker genes used for GTDB (bac120) classification (60,86) (Methods). Each of the largest 1,000 functions of our balanced organism set contained at least one protein associated with one of the 120 GTDB marker protein families. However, only slightly more than half (70 of the bac120) of the marker families were present in the 1,000 sets of 5,000 randomly selected Fusion functions. The remaining functions were most likely unique to individual organisms.

### Modularity-based taxonomic classification reflects phylogeny

Conventional taxonomic classification schemes rely on morphological and genetic markers (NCBI) or phylogenetic analysis of genetic data (GTDB). Genetic similarity, however, is not evenly spread across different sections of the taxonomy. Assuring that taxonomic groups at a given level are equally diverse is thus a well-known consideration when developing a taxonomy. GTDB, for example, tries to address this issue by breaking up the NCBI taxonomy’s polyphyletic taxa and reassigning organisms to taxonomic ranks higher than species in order to better represent genetic diversity at the individual level(86).

We clustered our organism functional similarity network, where organisms are vertices and edges represent Fusion functional similarity, computed based on the full set of functions, to extract groups of functionally related organisms – *Fusion-informed taxa* (Methods). We propose that this community detection-based taxonomy reflects functional similarity and metabolic/environmental preferences, and thus captures bacterial functional diversity better than phylogeny driven taxonomies. This is especially important when investigating environmentally specialized bacteria, e.g. symbionts or extremophiles, which are more likely to undergo convergent evolution and be functionally similar to other members of their environmental niche than to their phylogenetic relatives.

We identified resolution thresholds that influence the size and granularity of the Fusion-taxa such that the results best reflected existing taxonomic groupings at different taxonomic levels (Fig. 4). Note that for our balanced organism set, this excluded species and genus levels, as this set lacks pairs of organisms identical at these levels. We also note that this approach to threshold optimization was not meant to evaluate the ability of Fusion-based classification to precisely recall the existing taxonomy, but rather to identify plausible thresholds for augmenting the observable signal. To evaluate the similarity between Fusion-taxa and GTDB phylum/class/order/family levels we used the V-measure metric, using GTDB-taxon designations for organisms as reference labels and Fusion-taxa as predicted labels. The V-Measure is the harmonic mean between homogeneity and completeness; homogeneity is the number of organisms in a Fusion-taxon that belong to the same GTDB-taxon and completeness is the number of organisms of a GTDB-taxon that are found within one Fusion*-*taxon. A high V-measure indicates that both homogeneity and completeness are high. To delineate the Fusion-taxa from our organism network, we selected the Louvain(63) clustering resolutions attaining the highest V-measures (Fig. 4, Methods). For example, clustering the Fusion organism similarity network at a Louvain resolution =0.68 achieved the highest V-measure (among all other resolutions) when comparing the clusters’ members to the GTDB phyla members; that is, phylum and Fusion-taxon organisms coincided most at this resolution. Furthermore, distributions of GTDB taxon (phylum through order) sizes and the Fusion-taxon sizes (at the corresponding resolution) were similar; Kolmogorov-Smirnov *p*-values (values < 0.05 would indicate that two size distributions are different; Table S1). This observation suggests some similarity between the larger organism groups captured by Fusion and GTDB despite differences in their approaches to establishing organism relationships. However, the GTDB families were different in size from the corresponding Fusion-taxa (KS *p*-value = 0.01, Table S1), highlighting the (expected) divergence between the functional and phylogenetic approaches at finer taxonomic resolutions.

**Fig. 4.**
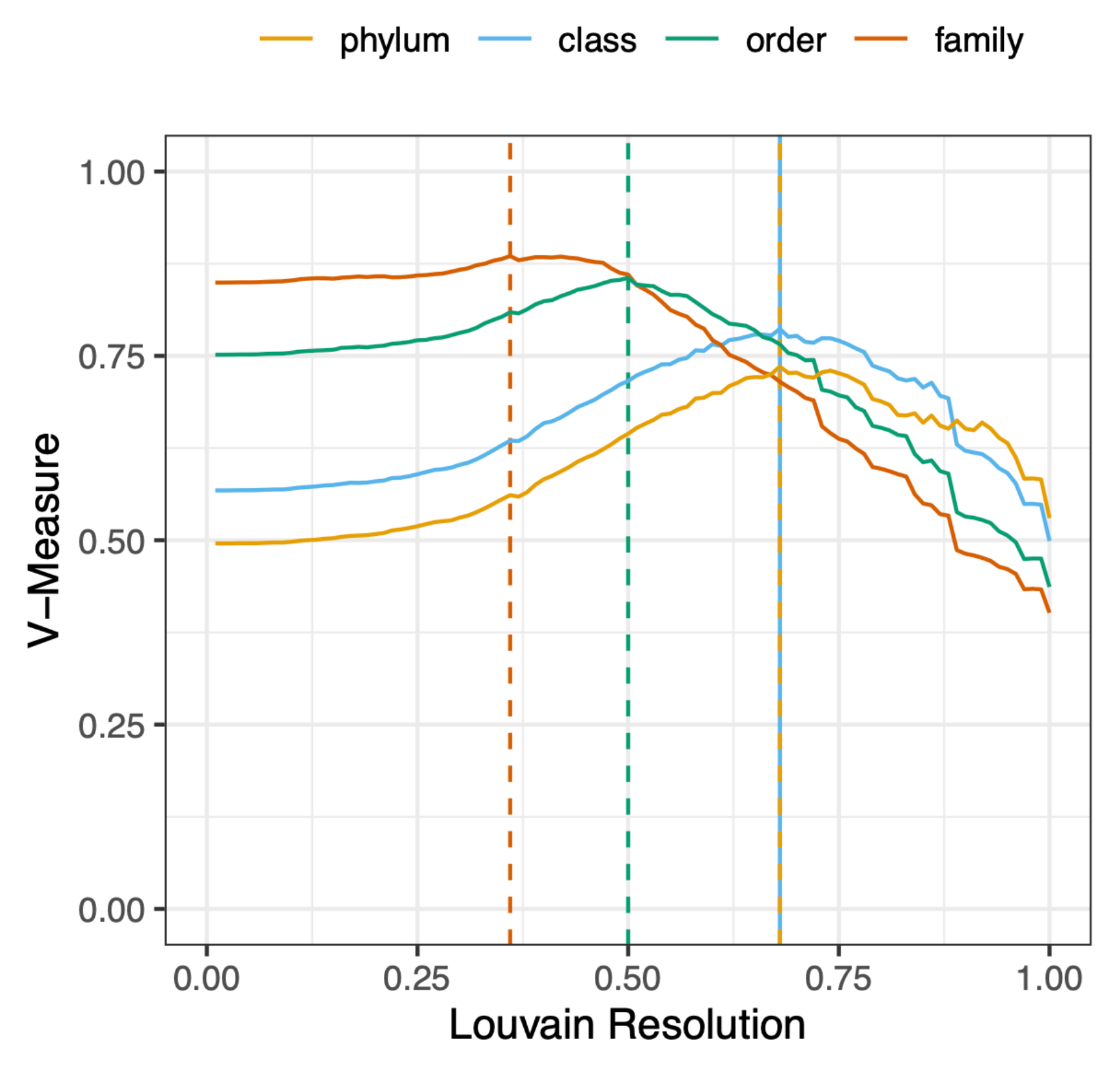
Community based organism classification using Fusion functional organism similarities recapitulates established taxonomy. Choosing different Louvain resolution parameters (x-axis) to establish communities of functionally similar organisms we can optimize the rate (y-axis) at which any two organisms are assigned to be in the same Fusion-taxon vs. reference of GTDB-taxonomy assignment. For example, clustering the Fusion organism similarity network at a Louvain resolution parameter of 0.36 yields the best approximation of communities of organisms, corresponding to the family taxonomic level. Thresholds for order, class and phylum are 0.50, 0.68 and 0.68 respectively.

### Modularity-based taxonomy is robust to the addition of novel organisms

As new organisms are added to taxonomies, organism assignments may need to be restructured. Here, updating the number of organisms per taxon or adding a new taxon containing only the novel organisms is far easier than reshuffling organisms from one taxon to others. Fusion-taxa appear robust to addition of organisms, favoring the former outcomes. To demonstrate this quality, we created 50,000 new organism similarity networks by adding *n* (*n* ∈ [1..500]) organisms to the balanced organism set clusters, i.e. 100 networks for each *n*, where *n* organisms not contained in the balanced set were randomly selected from the complete organism set; each network was of size of 1,503 to 2,002 organisms (balanced organisms set + *n*). We re-clustered all networks at resolution=0.5 (Methods), the resolution we previously determined to correspond best to the GTDB order-level classifications. The resulting clusters (predicted labels) of the balanced set organisms were compared to the original clusters (reference labels).

We expected that addition of these new organisms, selected from the complete set, and thus similar to those already in the network, would reflect the “worst case” scenario for network stability. That is, while new organisms could be expected to form their own clusters, microbes similar to those already in the network could stimulate cluster re-definition. Our function-based clustering did not change significantly upon addition of new (existing taxon) microbes, demonstrating the stability of the identified taxa (predicted vs. reference labels; with one added organism, median V-measure=0.99; with 500 added organisms: V-measure=0.96; Methods, Fig. S11).

To further evaluate the (likely limited) effects of introducing organisms of novel taxa, we extracted ten genomes added to GenBank after the date of our set extraction (February 2018) and whose GTDB order was not represented in our collection. We annotated the Fusion functional profiles of these organisms by running alignments, as in Zhu et al(33), against our set of proteins, computed organism similarities to the 1,502 microbes of our balanced set, and re-clustered the resulting network. Eight of these ten organisms each formed their own cluster, as expected. The two remaining organisms clustered into an already existing Fusion-taxon. Interestingly, this taxon contained an organism of the same NCBI order as the two new bacteria, illustrating the subjectivity of GTDB vs. NCBI taxonomies and highlighting the importance of organism assignment standardization.

Stability of our modularity-based taxonomy suggests that the functional space already covered by the current organism set is sufficiently large that adding a single novel organism is unlikely to require (the computationally expensive step of) reconfiguring the Fusion taxonomy. We expect that given a large enough number of novel organisms it may become sensible to regenerate the complete taxonomy. However, the currently observed levels of sequence and functional redundancy across the microbial space suggest that increasing the number of organisms should not drastically alter the Fusion taxonomy landscape in the near future.

### Machine learning-based sequence comparisons and sequence alignments capture different functional signals

Current function annotation methods are deeply steeped in function transfer by homology and often fail to annotate proteins with no obvious experimentally studied homologs (Prabakaran, R. and Bromberg, Y., Manuscript in Preparation). We built Fusion to be critically different from traditional classifiers and annotate putative (e.g. function cluster 45) instead of explicit (e.g. alcohol dehydrogenase) functionality. However, in our annotations we still heavily relied on significant sequence similarity. Thus, we aimed to build a function annotation model that was less dependent on sequence comparisons and somewhat arbitrary clustering parameters. To this end, we trained a Siamese Neural Network (SNN) to predict whether two nucleic acid (gene) sequences encode proteins of the same Fusion function.

SNNs are specifically optimized to assess similarities of two objects (87) – in our case gene/protein functional similarity. Note that we chose genes instead of protein sequences to adapt to unexplored genome and metagenome analyses more easily, where protein sequence prediction is an extra computational step in generating functional abundance profiles. In training (balanced set; ∼300K gene pairs, 50% same vs. 50% different function), our model attained 73% overall accuracy at the default cutoff (score>0.5; area under the ROC curve, AUC_ROC =0.80). SNN prediction scores correlated with the precision of recognizing the pair’s functional identity; thus, for example, at cutoff =0.98 the method attained 96% precision for the 19% of gene pairs that reached this threshold. Note that at this stringent cutoff, for an imbalanced test set with 10% same function pairs, the network still maintained high precision (82%) at a similar recall (24%). Importantly, increasing the size of the training data to one million gene pairs, improved the method performance (AUC_ROC = 0.81), suggesting that further improvements may be possible.

While somewhat correlated (Spearman rho=0.3, Fig. S12), the SNN similarity scores captured a different signal than the HFSP scores, i.e. values incorporating sequence identity and alignment length. To evaluate the captured signal, we compiled a set of Fusion functions where (1) the Fusion function was associated with only one EC number, (2) a number of different Fusion functions were associated with one EC number, and (3) different Fusion functions were associated with different EC numbers. As it was trained to do, SNN captured the similarity of genes from the first category (same Fusion function, same EC; Fig. S13 right green column, median SNN-score = 0.83) and the difference of the genes from the third category (different Fusion function, different EC; median SNN-score = 0.13; Fig. S13, left orange column). However, genes of the second category (different Fusion functions, same EC number) were scored significantly higher than expected (median SNN-score = 0.7; Fig. S13, left green column; false positives in SNN training.) Thus, our SNN identified same enzymatic activity gene pairs that were NOT captured as same function by the homology-based Fusion.

### Machine learning-based sequence comparisons and structure alignments capture orthogonal signals

What functional similarity does an SNN capture? While not obvious from gene sequence comparisons, we expected that functionally similar proteins that are not sequence similar should share structural similarity(88,89). We compiled a set of Fusion proteins that have a structure in the PDB and then computed structural (TM-scores) and functional (SNN-scores) similarities for all pairs (Methods). Note that we did not use predicted protein structures(90,91) to avoid compounding machine learning preferences.

First, we examined the relationship between the TM-score and SNN-score for sequence-similar protein pairs (HFSP ≥0; Fig. S14). We found that 97% of these pairs (3,988 of 4,132) were structurally similar (TM-score ≥0.7; Table S2) and 94% (3,876) were predicted by the SNN to be of the same function (SNN-score ≥0.5; Table S3). These observations highlight HFSP’s precision and confirm the expectation that high sequence similarity in most cases encodes for structural and functional identity. Note that only a fifth (3,988 of 17,872) of all protein pairs with a TM-score ≥0.7 also had an HFSP≥0, indicating that function transfer by homology, while precise for the pairs it does identify, fails to find the more remote functional similarity.

SNN predictions, on the other hand, identified 77% (13,750 of 17,862) of the high TM-scoring pairs to be of the same function. Note that a quarter (3,583 of 13,750) of the SNN predictions attained a high score (≥0.98; Fig. 5, Table S4) but only some of these (2,119; 59%) were also sequence similar (HFSP≥0). Most (73%, 21,668 of 29,531) of the reliably structurally dissimilar protein pairs (TM-scores <0.2, Methods) were predicted to be functionally different by SNN (score <0.5); only 81 pairs (<1%) attained a high SNN-score. Of pairs that share some structural similarity, SNN labeled half (TM-score: [0.2,0.5) =45% and [0.5,0.7)=53%) as having the same Fusion function; for both sets, only 4% reached a high SNN-score, which stands in contrast to the 26% of the protein pairs with TM-score ≥0.7. That is, SNN reliably identifies presence/absence of functional similarity at the extremes of structural similarity but is significantly less certain for mildly structurally similar proteins.

**Fig. 5.**
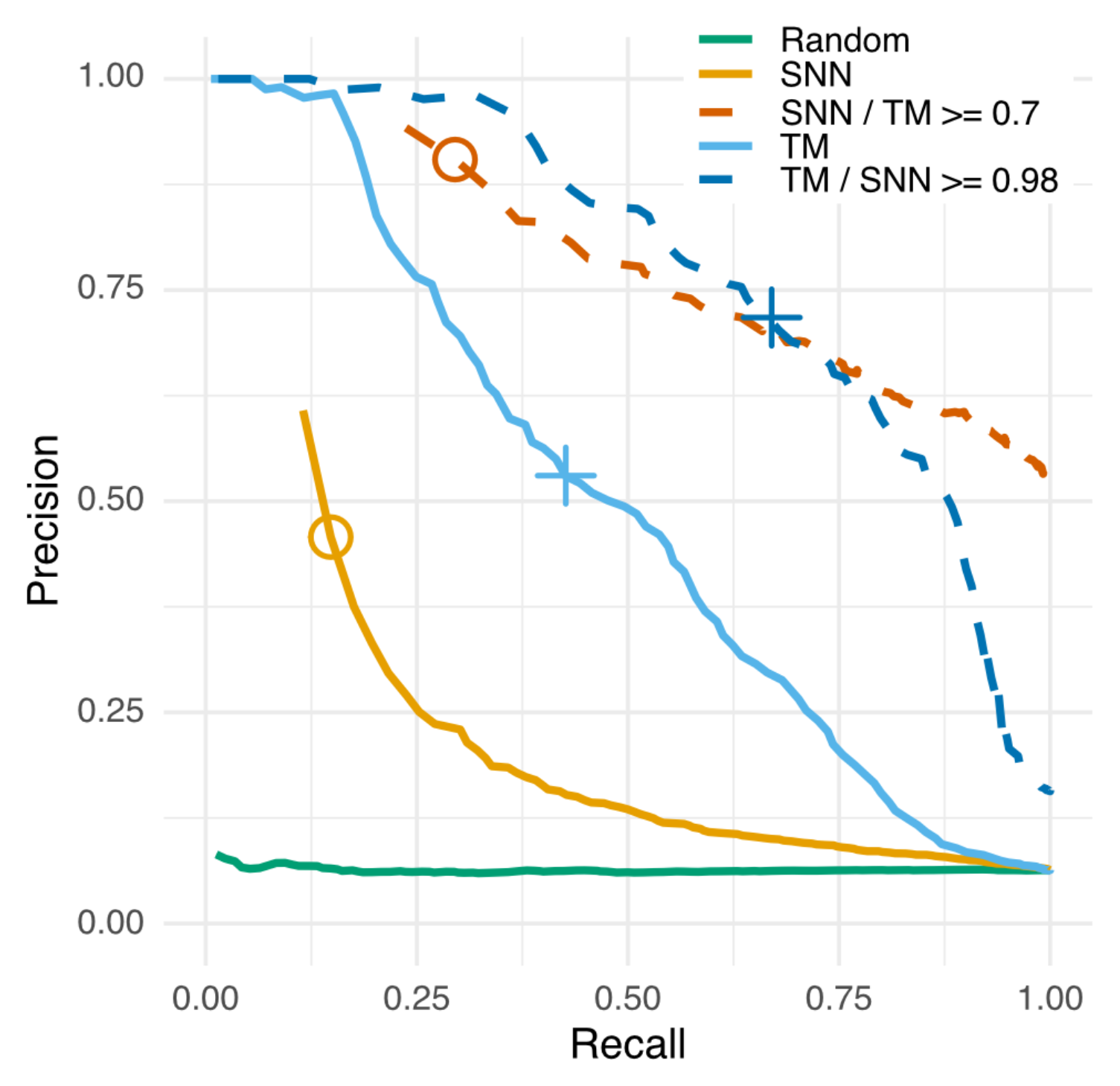
Combining TM and SNN scores improves annotation of functionally similar proteins. For proteins with available structures, the TM-score (blue solid line) was a better estimate of protein functional similarity (same EC number) than the SNN-score (orange solid line); even at the high reliability threshold of SNN-score ≥0.98 (circle), the method attained only 46% precision and 16% recall as compared 53% precision and 43% recall of the TM-score≥0.7 (cross). However, the combined SNN & TM-score metrics (dashed lines) were better than either of the methods alone. That is, for a subset of structurally similar proteins (TM≥0.7) the SNN score (orange dashed line) was a good indicator of functional similarity. Similarly, for reliably functionally similar proteins (SNN≥0.98), the TM-score (blue dashed line) had a significantly higher precision. Note that our dataset is representative of real life and thus, trivially, imbalanced as there are significantly fewer same EC (positive) pairs than different EC (negative) pairs; here, a ratio of ∼1/15.

We further evaluated the ability of the SNN and the TM scores to directly predict function, i.e. the identify/difference of the experimentally annotated 3^rd^ EC level of each protein pair (Methods). As before (Fig. S13), we observed that the proteins of the same EC number were, on average, predicted with a higher SNN-score than different-EC pairs (Fig. S15). We found that while the SNN precision and recall were significantly above random, they were lower than simply using the TM-score (Fig. 5). However, combining the TM and SNN predictions significantly improved recognition of proteins of the same function. We thus suggest that the SNN reports a signal of functional similarity that is captured neither by sequence nor structure similarity alone.

To explore this signal further, we investigated outlier protein pairs in our set, i.e. structurally different (TM-score <0.2), sequence dissimilar (HFSP<0) pairs of proteins of the same 4^th^ digit EC number attaining an SNNscore≥0.98, i.e. UniProt ids: P37870/P37871, O35011/O31718, and Q8RQE9 /P3787. Curiously, for these pairs, both TMAlign (proteins in the pair have different structures) and the SNN (proteins in the pair have the same function) predictions were correct. That is, each pair was annotated with one EC number, but the sequences were structurally different chains of the same heteromer (P37870/P37871 and O35011/O31718) or of the same protein complex (Q8RQE9/P37871). While these three examples are anecdotal evidence they also clearly demonstrate the limitations of available chain-based functional annotations.

### Summarizing the Findings

Understanding bacterial lifestyles requires describing their functional capabilities and critically contributes to research in medical, environmental, and industrial fields. The recent explosion in completely sequenced bacterial genomes has, simultaneously, created a deluge of functionally unannotated and misannotated sequences and allowed for the development of new and informative sequence-based methods. Here, we optimized Fusion, a method for annotating the functional repertoires of bacteria. Using these repertoires, we propose an implementation of a function-based taxonomy that is robust to addition of new organisms. We note that Fusion functions recapitulate bacterial taxonomic assignments better than 16S rRNA comparisons. In the absence of validation techniques for functional assignment of unannotated proteins, we suggest that taxonomic classification abilities highlight Fusion’s precision in capturing putative protein functionality. In further exploration of the functional signal, we trained a Siamese Neural Network (SNN) to label pairs of genes whose product proteins are functionally similar according to our functional definitions. Notably, the SNN’s predictions were orthogonal to sequence and structure signals and, thus, may open the door to investigating remote homology. We propose that this method could further be optimized for extraction of functional annotation directly from metagenomic reads.

## AVAILABILITY

All data are available in the main text, the Supplementary Materials or referenced permanent online data repositories: Function dataset: 10.6084/m9.figshare.21599544, Organism similarities : 10.6084/m9.figshare.21637988, Structural similarities: 10.6084/m9.figshare.21637937, Jupyter & R notebooks containing the analysis of datasets: https://bitbucket.org/bromberglab/fusion-manuscript-analysis/, git-repository containing the code to generate the Fusion functions: https://bitbucket.org/bromberglab/fusion-updater/, git-repository containing code to infer functional similarity between organisms: https://bitbucket.org/bromberglab/fusion-organism_sim/, git-repository containing code to evaluate functional similarity of proteins based on Fold-seek & our SNN: https://bitbucket.org/bromberglab/fusion-snn/

## ACKNOWLEDGEMENT

We want to thank Ariel Aptekmann, Maximilian Miller, Zishou Zeng, and Peter Kahn (all Rutgers University) for the valuable feedback during the concept phase of the project and Prabakaran Ramakrishnan (Emory University) for critical reading of the manuscript. Many thanks to Martin Steinegger (Seoul National University) for assistance with MMSeqs2 and Foldseek. Lastly, we extend our gratitude to everyone in the scientific community for providing the tools, databases and data sources that were vital in producing this research.

## REFERENCES

1. Blaser, M.J., Cardon, Z.G., Cho, M.K., Dangl, J.L., Donohue, T.J., Green, J.L., Knight, R., Maxon, M.E., Northen, T.R., Pollard, K.S. et al. (2016) Toward a Predictive Understanding of Earth&#x2019;s Microbiomes to Address 21st Century Challenges. mBio, 7, e00714–00716.

2. Falkowski, P.G., Fenchel, T. and Delong, E.F. (2008) The Microbial Engines That Drive Earth’s Biogeochemical Cycles. Science, 320, 1034–1039.

3. Jousset, A., Bienhold, C., Chatzinotas, A., Gallien, L., Gobet, A., Kurm, V., Küsel, K., Rillig, M.C., Rivett, D.W., Salles, J.F. et al. (2017) Where less may be more: how the rare biosphere pulls ecosystems strings. The ISME Journal, 11, 853–862.

4. Russell, J.A., Dubilier, N. and Rudgers, J.A. (2014) Nature’s microbiome: introduction. Molecular Ecology, 23, 1225–1237.

5. Bromberg, Y., Aptekmann, A.A., Mahlich, Y., Cook, L., Senn, S., Miller, M., Nanda, V., Ferreiro, D.U. and Falkowski, P.G. (2022) Quantifying structural relationships of metal-binding sites suggests origins of biological electron transfer. Science Advances, 8, eabj3984.

6. Shade, A. (2018) Understanding Microbiome Stability in a Changing World. mSystems, 3.

7. Beghini, F., McIver, L.J., Blanco-Míguez, A., Dubois, L., Asnicar, F., Maharjan, S., Mailyan, A., Manghi, P., Scholz, M., Thomas, A.M. et al. (2021) Integrating taxonomic, functional, and strain-level profiling of diverse microbial communities with bioBakery 3. eLife, 10, e65088.

8. Franzosa, E.A., McIver, L.J., Rahnavard, G., Thompson, L.R., Schirmer, M., Weingart, G., Lipson, K.S., Knight, R., Caporaso, J.G., Segata, N. et al. (2018) Species-level functional profiling of metagenomes and metatranscriptomes. Nature Methods, 15, 962–968.

9. Kaminski, J., Gibson, M.K., Franzosa, E.A., Segata, N., Dantas, G. and Huttenhower, C. (2015) High-Specificity Targeted Functional Profiling in Microbial Communities with ShortBRED. PLOS Computational Biology, 11, e1004557.

10. Bolyen, E., Rideout, J.R., Dillon, M.R., Bokulich, N.A., Abnet, C.C., Al-Ghalith, G.A., Alexander, H., Alm, E.J., Arumugam, M., Asnicar, F. et al. (2019) Reproducible, interactive, scalable and extensible microbiome data science using QIIME 2. Nature Biotechnology, 37, 852–857.

11. Stackebrandt, E. and Goebel, B.M. (1994) Taxonomic note: a place for DNA-DNA reassociation and 16S rRNA sequence analysis in the present species definition in bacteriology. International journal of systematic and evolutionary microbiology, 44, 846–849.

12. Brenner, D.J., Fanning, G.R., Johnson, K.E., Citarella, R.V. and Falkow, S. (1969) Polynucleotide sequence relationships among members of Enterobacteriaceae. J Bacteriol, 98, 637–650.

13. Goris, J., Konstantinidis, K.T., Klappenbach, J.A., Coenye, T., Vandamme, P. and Tiedje, J.M. (2007) DNA-DNA hybridization values and their relationship to whole-genome sequence similarities. Int J Syst Evol Microbiol, 57, 81–91.

14. Boone, D.R., Castenholz, R.W., Garrity, G.M. and Stanley, J. (2001) Bergey’s Manual® of Systematic Bacteriology: Volume One The Archaea and the Deeply Branching and Phototrophic Bacteria. Springer.

15. Woese, C.R., Kandler, O. and Wheelis, M.L. (1990) Towards a natural system of organisms: proposal for the domains Archaea, Bacteria, and Eucarya. Proceedings of the National Academy of Sciences, 87, 4576–4579.

16. Johnson, J.S., Spakowicz, D.J., Hong, B.-Y., Petersen, L.M., Demkowicz, P., Chen, L., Leopold, S.R., Hanson, B.M., Agresta, H.O., Gerstein, M. et al. (2019) Evaluation of 16S rRNA gene sequencing for species and strain-level microbiome analysis. Nature Communications, 10, 5029.

17. Konstantinidis, K.T., Ramette, A. and Tiedje, J.M. (2006) Toward a More Robust Assessment of Intraspecies Diversity, Using Fewer Genetic Markers. Applied and Environmental Microbiology, 72, 7286–7293.

18. Konstantinidis, K.T. and Tiedje, J.M. (2007) Prokaryotic taxonomy and phylogeny in the genomic era: advancements and challenges ahead. Current Opinion in Microbiology, 10, 504–509.

19. Gevers, D., Cohan, F.M., Lawrence, J.G., Spratt, B.G., Coenye, T., Feil, E.J., Stackebrandt, E., de Peer, Y.V., Vandamme, P., Thompson, F.L. et al. (2005) Re-evaluating prokaryotic species. Nature Reviews Microbiology, 3, 733–739.

20. Rosselló-Mora, R. (2005) Updating Prokaryotic Taxonomy. Journal of Bacteriology, 187, 6255–6257.

21. Gevers, D., Dawyndt, P., Vandamme, P., Willems, A., Vancanneyt, M., Swings, J. and De Vos, P. (2006) Stepping stones towards a new prokaryotic taxonomy. Philosophical Transactions of the Royal Society B: Biological Sciences, 361, 1911–1916.

22. Hilario, E. and Gogarten, J.P. (1993) Horizontal transfer of ATPase genes — the tree of life becomes a net of life. Biosystems, 31, 111–119.

23. Babić, A., Lindner, A.B., Vulić, M., Stewart, E.J. and Radman, M. (2008) Direct Visualization of Horizontal Gene Transfer. Science, 319, 1533–1536.

24. Goldenfeld, N. and Woese, C. (2007) Biology’s next revolution. Nature, 445, 369–369.

25. Price, M.N., Dehal, P.S. and Arkin, A.P. (2008) Horizontal gene transfer and the evolution of transcriptional regulation in Escherichia coli. Genome Biology, 9, R4.

26. He, Z., Xu, S. and Shi, S. (2020) Adaptive convergence at the genomic level—prevalent, uncommon or very rare? National Science Review, 7, 947–951.

27. Farhat, M.R., Shapiro, B.J., Kieser, K.J., Sultana, R., Jacobson, K.R., Victor, T.C., Warren, R.M., Streicher, E.M., Calver, A., Sloutsky, A. et al. (2013) Genomic analysis identifies targets of convergent positive selection in drug-resistant Mycobacterium tuberculosis. Nature Genetics, 45, 1183–1189.

28. Zhu, C., Delmont, T.O., Vogel, T.M. and Bromberg, Y. (2015) Functional basis of microorganism classification. PLoS Comput Biol, 11, e1004472.

29. Rastogi, G. and Sani, R.K. (2011) In Ahmad, I., Ahmad, F. and Pichtel, J. (eds.), Microbes and Microbial Technology: Agricultural and Environmental Applications. Springer New York, New York, NY, pp. 29–57.

30. Langille, M.G.I., Zaneveld, J., Caporaso, J.G., McDonald, D., Knights, D., Reyes, J.A., Clemente, J.C., Burkepile, D.E., Vega Thurber, R.L., Knight, R. et al. (2013) Predictive functional profiling of microbial communities using 16S rRNA marker gene sequences. Nature Biotechnology, 31, 814–821.

31. Schleifer, K.H. (2009) Classification of Bacteria and Archaea: Past, present and future. Systematic and Applied Microbiology, 32, 533–542.

32. Young, J.M. (2001) Implications of alternative classifications and horizontal gene transfer for bacterial taxonomy. International Journal of Systematic and Evolutionary Microbiology, 51, 945–953.

33. Zhu, C., Mahlich, Y., Miller, M. and Bromberg, Y. (2018) Fusion DB: assessing microbial diversity and environmental preferences via functional similarity networks. Nucleic acids research, 46, D535–D541.

34. Bromley, J., Guyon, I., LeCun, Y., Säckinger, E. and Shah, R. (1993) Signature verification using a" siamese" time delay neural network. Advances in neural information processing systems, 6.

35. Koehler Leman, J., Szczerbiak, P., Renfrew, P.D., Gligorijevic, V., Berenberg, D., Vatanen, T., Taylor, B.C., Chandler, C., Janssen, S., Pataki, A., et al. (2023) Sequence-structure-function relationships in the microbial protein universe. Nature Communications, 14, 2351.

36. Nissen, J.N., Johansen, J., Allesøe, R.L., Sønderby, C.K., Armenteros, J.J.A., Grønbech, C.H., Jensen, L.J., Nielsen, H.B., Petersen, T.N., Winther, O. et al. (2021) Improved metagenome binning and assembly using deep variational autoencoders. Nature Biotechnology, 39, 555–560.

37. Pan, S., Zhu, C., Zhao, X.-M. and Coelho, L.P. (2022) A deep siamese neural network improves metagenome-assembled genomes in microbiome datasets across different environments. Nature Communications, 13, 2326.

38. Kang, D.D., Li, F., Kirton, E., Thomas, A., Egan, R., An, H. and Wang, Z. (2019) MetaBAT 2: an adaptive binning algorithm for robust and efficient genome reconstruction from metagenome assemblies. PeerJ, 7, e7359.

39. Wu, Y.-W., Simmons, B.A. and Singer, S.W. (2015) MaxBin 2.0: an automated binning algorithm to recover genomes from multiple metagenomic datasets. Bioinformatics, 32, 605–607.

40. Benson, D.A., Cavanaugh, M., Clark, K., Karsch-Mizrachi, I., Lipman, D.J., Ostell, J. and Sayers, E.W. (2013) GenBank. Nucleic Acids Res, 41, D36–42.

41. Sayers, E.W., Cavanaugh, M., Clark, K., Ostell, J., Pruitt, K.D. and Karsch-Mizrachi, I. (2019) GenBank. Nucleic Acids Res, 47, D94–d99.

42. Li, W. and Godzik, A. (2006) Cd-hit: a fast program for clustering and comparing large sets of protein or nucleotide sequences. Bioinformatics, 22, 1658–1659.

43. Fu, L., Niu, B., Zhu, Z., Wu, S. and Li, W. (2012) CD-HIT: accelerated for clustering the next-generation sequencing data. Bioinformatics, 28, 3150–3152.

44. Mahlich, Y., Steinegger, M., Rost, B. and Bromberg, Y. (2018) HFSP: high speed homology-driven function annotation of proteins. Bioinformatics, 34, i304–i312.

45. Steinegger, M. and Söding, J. (2017) MMseqs2 enables sensitive protein sequence searching for the analysis of massive data sets. Nat Biotechnol, 35, 1026–1028.

46. Azad, A., Pavlopoulos, G.A., Ouzounis, C.A., Kyrpides, N.C. and Buluç, A. (2018) HipMCL: a high-performance parallel implementation of the Markov clustering algorithm for large-scale networks. Nucleic Acids Res, 46, e33.

47. Van Dongen, S.M. (2000).

48. Enright, A.J., Van Dongen, S. and Ouzounis, C.A. (2002) An efficient algorithm for large-scale detection of protein families. Nucleic Acids Res, 30, 1575–1584.

49. Bairoch, A. (2000) The ENZYME database in 2000. Nucleic Acids Research, 28, 304–305.

50. Bairoch, A. and Apweiler, R. (1997) The SWISS-PROT protein sequence database: its relevance to human molecular medical research. J Mol Med (Berl), 75, 312–316.

51. Consortium, T.U. (2020) UniProt: the universal protein knowledgebase in 2021. Nucleic Acids Research, 49, D480–D489.

52. El-Gebali, S., Mistry, J., Bateman, A., Eddy, S.R., Luciani, A., Potter, S.C., Qureshi, M., Richardson, L.J., Salazar, G.A., Smart, A. et al. (2018) The Pfam protein families database in 2019. Nucleic Acids Research, 47, D427–D432.

53. Madeira, F., Park, Y.M., Lee, J., Buso, N., Gur, T., Madhusoodanan, N., Basutkar, P., Tivey, A.R., Potter, S.C. and Finn, R.D. (2019) The EMBL-EBI search and sequence analysis tools APIs in 2019. Nucleic acids research, 47, W636–W641.

54. Eddy, S.R. (2011) Accelerated Profile HMM Searches. PLOS Computational Biology, 7, e1002195.

55. (2021) The Gene Ontology resource: enriching a GOld mine. Nucleic Acids Research, 49, D325–D334.

56. Ashburner, M., Ball, C.A., Blake, J.A., Botstein, D., Butler, H., Cherry, J.M., Davis, A.P., Dolinski, K., Dwight, S.S., Eppig, J.T. et al. (2000) Gene ontology: tool for the unification of biology. The Gene Ontology Consortium. Nat Genet, 25, 25–29.

57. Rosenberg, A. and Hirschberg, J. (2007), Proceedings of the 2007 joint conference on empirical methods in natural language processing and computational natural language learning (EMNLP-CoNLL), pp. 410–420.

58. Pedregosa, F., Varoquaux, G., Gramfort, A., Michel, V., Thirion, B., Grisel, O., Blondel, M., Prettenhofer, P., Weiss, R. and Dubourg, V. (2011) Scikit-learn: Machine learning in Python. the Journal of machine Learning research, 12, 2825–2830.

59. Schoch, C.L., Ciufo, S., Domrachev, M., Hotton, C.L., Kannan, S., Khovanskaya, R., Leipe, D., McVeigh, R., O’Neill, K., Robbertse, B. et al. (2020) NCBI Taxonomy: a comprehensive update on curation, resources and tools. Database (Oxford), 2020.

60. Parks, D.H., Chuvochina, M., Chaumeil, P.-A., Rinke, C., Mussig, A.J. and Hugenholtz, P. (2020) A complete domain-to-species taxonomy for Bacteria and Archaea. Nature Biotechnology, 1-8.

61. Sukumaran, J. and Holder, M.T. (2010) DendroPy: a Python library for phylogenetic computing. Bioinformatics, 26, 1569–1571.

62. Menardo, F., Loiseau, C., Brites, D., Coscolla, M., Gygli, S.M., Rutaihwa, L.K., Trauner, A., Beisel, C., Borrell, S. and Gagneux, S. (2018) Treemmer: a tool to reduce large phylogenetic datasets with minimal loss of diversity. BMC Bioinformatics, 19, 164.

63. Blondel, V.D., Guillaume, J.-L., Lambiotte, R. and Lefebvre, E. (2008) Fast unfolding of communities in large networks. Journal of statistical mechanics: theory and experiment, 2008, P10008.

64. Cole, J.R., Wang, Q., Fish, J.A., Chai, B., McGarrell, D.M., Sun, Y., Brown, C.T., Porras-Alfaro, A., Kuske, C.R. and Tiedje, J.M. (2014) Ribosomal Database Project: data and tools for high throughput rRNA analysis. Nucleic Acids Res, 42, D633–642.

65. Nawrocki, E.P. and Eddy, S.R. (2013) Infernal 1.1: 100-fold faster RNA homology searches. Bioinformatics, 29, 2933–2935.

66. Ondov, B.D., Treangen, T.J., Melsted, P., Mallonee, A.B., Bergman, N.H., Koren, S. and Phillippy, A.M. (2016) Mash: fast genome and metagenome distance estimation using MinHash. Genome Biology, 17, 132.

67. Hoarfrost, A., Aptekmann, A., Farfanuk, G. and Bromberg, Y. (2020) Shedding Light on Microbial Dark Matter with A Universal Language of Life. bioRxiv.

68. Hochreiter, S. and Schmidhuber, J. (1997) Long Short-Term Memory. Neural Computation, 9, 1735–1780.

69. Burley, S.K., Bhikadiya, C., Bi, C., Bittrich, S., Chen, L., Crichlow, G.V., Christie, C.H., Dalenberg, K., Di Costanzo, L., Duarte, J.M. et al. (2020) RCSB Protein Data Bank: powerful new tools for exploring 3D structures of biological macromolecules for basic and applied research and education in fundamental biology, biomedicine, biotechnology, bioengineering and energy sciences. Nucleic Acids Research, 49, D437–D451.

70. Berman, H.M., Westbrook, J., Feng, Z., Gilliland, G., Bhat, T.N., Weissig, H., Shindyalov, I.N. and Bourne, P.E. (2000) The Protein Data Bank. Nucleic Acids Research, 28, 235–242.

71. van Kempen, M., Kim, S.S., Tumescheit, C., Mirdita, M., Söding, J. and Steinegger, M. (2022) Foldseek: fast and accurate protein structure search. bioRxiv, 2022.2002.2007.479398.

72. Zhang, Y. and Skolnick, J. (2004) Scoring function for automated assessment of protein structure template quality. *Proteins: Structure*, Function, and Bioinformatics, 57, 702–710.

73. Zhang, Y. and Skolnick, J. (2005) TM-align: a protein structure alignment algorithm based on the TM-score. Nucleic acids research, 33, 2302–2309.

74. Mistry, J., Chuguransky, S., Williams, L., Qureshi, M., Salazar, Gustavo A., Sonnhammer, E.L.L., Tosatto, S.C.E., Paladin, L., Raj, S., Richardson, L.J. et al. (2020) Pfam: The protein families database in 2021. Nucleic Acids Research, 49, D412–D419.

75. Barrera, A., Alastruey-Izquierdo, A., Martín, M.J., Cuesta, I. and Vizcaíno, J.A. (2014) Analysis of the Protein Domain and Domain Architecture Content in Fungi and Its Application in the Search of New Antifungal Targets. PLOS Computational Biology, 10, e1003733.

76. Koonin, E.V., Wolf, Y.I. and Karev, G.P. (2002) The structure of the protein universe and genome evolution. Nature, 420, 218–223.

77. Itoh, M., Nacher, J.C., Kuma, K.-i., Goto, S. and Kanehisa, M. (2007) Evolutionary history and functional implications of protein domains and their combinations in eukaryotes. Genome Biology, 8, R121.

78. Peisajovich, S.G., Garbarino, J.E., Wei, P. and Lim, W.A. (2010) Rapid Diversification of Cell Signaling Phenotypes by Modular Domain Recombination. Science, 328, 368–372.

79. Radivojac, P. (2022) Advancing remote homology detection: A step toward understanding and accurately predicting protein function. Cell Syst, 13, 435–437.

80. Rosselló-Mora, R. and Amann, R. (2001) The species concept for prokaryotes. FEMS Microbiology Reviews, 25, 39–67.

81. Větrovský, T. and Baldrian, P. (2013) The variability of the 16S rRNA gene in bacterial genomes and its consequences for bacterial community analyses. PLoS One, 8, e57923.

82. Zhu, C., Miller, M., Marpaka, S., Vaysberg, P., Rühlemann, M.C., Wu, G., Heinsen, F.-A., Tempel, M., Zhao, L., Lieb, W. et al. (2017) Functional sequencing read annotation for high precision microbiome analysis. Nucleic Acids Research, 46, e23–e23.

83. Hernández-Salmerón, J.E. and Moreno-Hagelsieb, G. (2022) FastANI, Mash and Dashing equally differentiate between Klebsiella species. PeerJ, 10, e13784.

84. Jain, C., Rodriguez-R, L.M., Phillippy, A.M., Konstantinidis, K.T. and Aluru, S. (2018) High throughput ANI analysis of 90K prokaryotic genomes reveals clear species boundaries. Nature Communications, 9, 5114.

85. Baker, D.N. and Langmead, B. (2019) Dashing: fast and accurate genomic distances with HyperLogLog. Genome Biol, 20, 265.

86. Parks, D.H., Chuvochina, M., Waite, D.W., Rinke, C., Skarshewski, A., Chaumeil, P.-A. and Hugenholtz, P. (2018) A standardized bacterial taxonomy based on genome phylogeny substantially revises the tree of life. Nature Biotechnology, 36, 996–1004.

87. Chicco, D. (2021) In Cartwright, H. (ed.), Artificial Neural Networks. Springer US, New York, NY, pp. 73–94.

88. Krissinel, E. (2007) On the relationship between sequence and structure similarities in proteomics. Bioinformatics, 23, 717–723.

89. Rost, B. (1999) Twilight zone of protein sequence alignments. *Protein Engineering*, Design and Selection, 12, 85–94.

90. Jumper, J., Evans, R., Pritzel, A., Green, T., Figurnov, M., Ronneberger, O., Tunyasuvunakool, K., Bates, R., Žídek, A., Potapenko, A., et al. (2021) Highly accurate protein structure prediction with AlphaFold. Nature, 596, 583–589.

91. Baek, M., DiMaio, F., Anishchenko, I., Dauparas, J., Ovchinnikov, S., Lee, G.R., Wang, J., Cong, Q., Kinch, L.N., Schaeffer, R.D. et al. (2021) Accurate prediction of protein structures and interactions using a three-track neural network. Science, 373, 871–876.

