## Supplementary for "Learning from the unknown: exploring the range of bacterial functionality"

**This PDF file includes:**

Supplementary Methods
Supplementary Results
Figs. S1 to S15
Tables S1 to S4

**SUPPLEMENTARY METHODS**

(Eqn. S1)

$$h = \begin{cases} 1 & if\ H(C,K) = 0 \\ 1 - \frac{H(C|K)}{H(C)} & else \end{cases}$$ where

$$H(C|K) = -\sum_{k=1}^{|K|}\sum_{c=1}^{|C|}\frac{a_{ck}}{N}\,log\frac{a_{ck}}{\sum_{c=1}^{|C|} a_{ck}}$$

$$H(C) = -\sum_{c=1}^{|C|}\frac{\sum_{k=1}^{|K|} a_{ck}}{n}\,log\frac{\sum_{k=1}^{|K|} a_{ck}}{n}$$

Where *N* the number of data points, *C* is the set of reference classes, *K* is the set of predicted cluster labels. *n* is the total number of references classes *c*. $a_{ck}$ is the number of data points that are both in reference class *c* and predicted cluster *k*.

(Eqn. S2) $$c = \begin{cases} 1 & if\ H(K,C) = 0 \\ 1 - \frac{H(K|C)}{H(K)} & else \end{cases}$$ where

$$H(K|C) = -\sum_{c=1}^{|C|}\sum_{k=1}^{|K|}\frac{a_{ck}}{N}\,log\frac{a_{ck}}{\sum_{k=1}^{|K|} a_{ck}}$$

$$H(K) = -\sum_{k=1}^{|K|}\frac{\sum_{c=1}^{|C|} a_{ck}}{n}\,log\frac{\sum_{c=1}^{|C|} a_{ck}}{n}$$

(Eqn. S3) $v = \frac{2*h*c}{h+c}$

### SUPPLEMENTARY Results

**Sequenced bacterial proteomes are significantly redundant.** We retrieved from GenBank (*1, 2*) (Methods) a set of 8,906 genomes/proteomes of bacterial organisms representing 3,005 species. This set comprised all fully sequenced bacterial genomes available at the time of extraction (2018). It is notably redundant with 65% (5,754) of the proteomes belonging to only 25% (753) of the species. An extreme case of this observation is the 360 proteomes of *Bordetella pertussis,* contributing 360 copies of almost every *B.pertussis* protein (~3,620 proteins per proteome) to our collection. Overall, nearly 60% (18.8 of 31.6 million) of proteins in our set were identical to others. Of the ~15.6M sequence-unique proteins in our set (*sequence-unique protein set*, Methods), ~2.8M (~18%) were found in multiple proteomes, while ~12.8M (82%) were proteome-specific. Similar observations have been reported previously (*3*).

As expected, much of the sequence redundancy occurred between strains of the same species, emphasizing the difficulty of distinguishing organism classes. When the set of organisms was phylogenetically balanced (*balanced organism set*, Methods), much of the protein redundancy was removed. This collection contained ~4.75 million proteins, of which 99% (~4.69M) were sequence- and organism- unique. We also note that these proteins still recapitulated nearly two thirds of the functions identified in the complete set of proteins (Methods). Most of the analyses presented here are based on the balanced organism set.

**Fig. S1.**

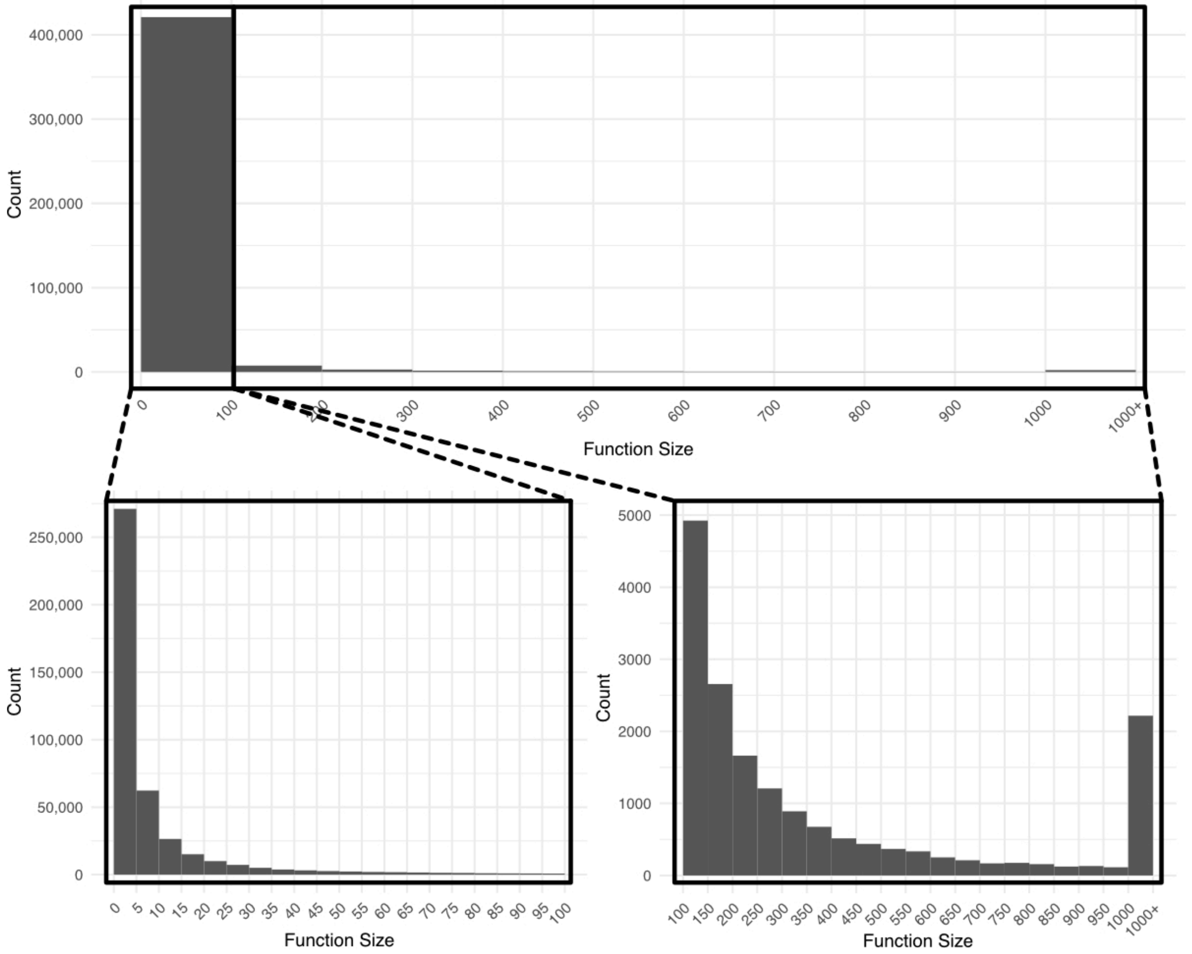


**Fig. S1. Nearly all fusion functions contain less than 100 proteins.** The top panel shows the distribution of function sizes binned by sizes of 100 (e.g. the second bin contains all functions of size 101-200). The last bin counts all functions of size > 1000. Functions of size <= 100 are further split into a distribution of bin size 5 (bottom left panel) and functions of size > 100 into bins of 50 (bottom right panel). More than 50% of all functions (~260k of ~440k) are of size 5 or smaller. On the other hand only a little of 2000 functions contain more than 1,000 proteins.

**Fig. S2.**

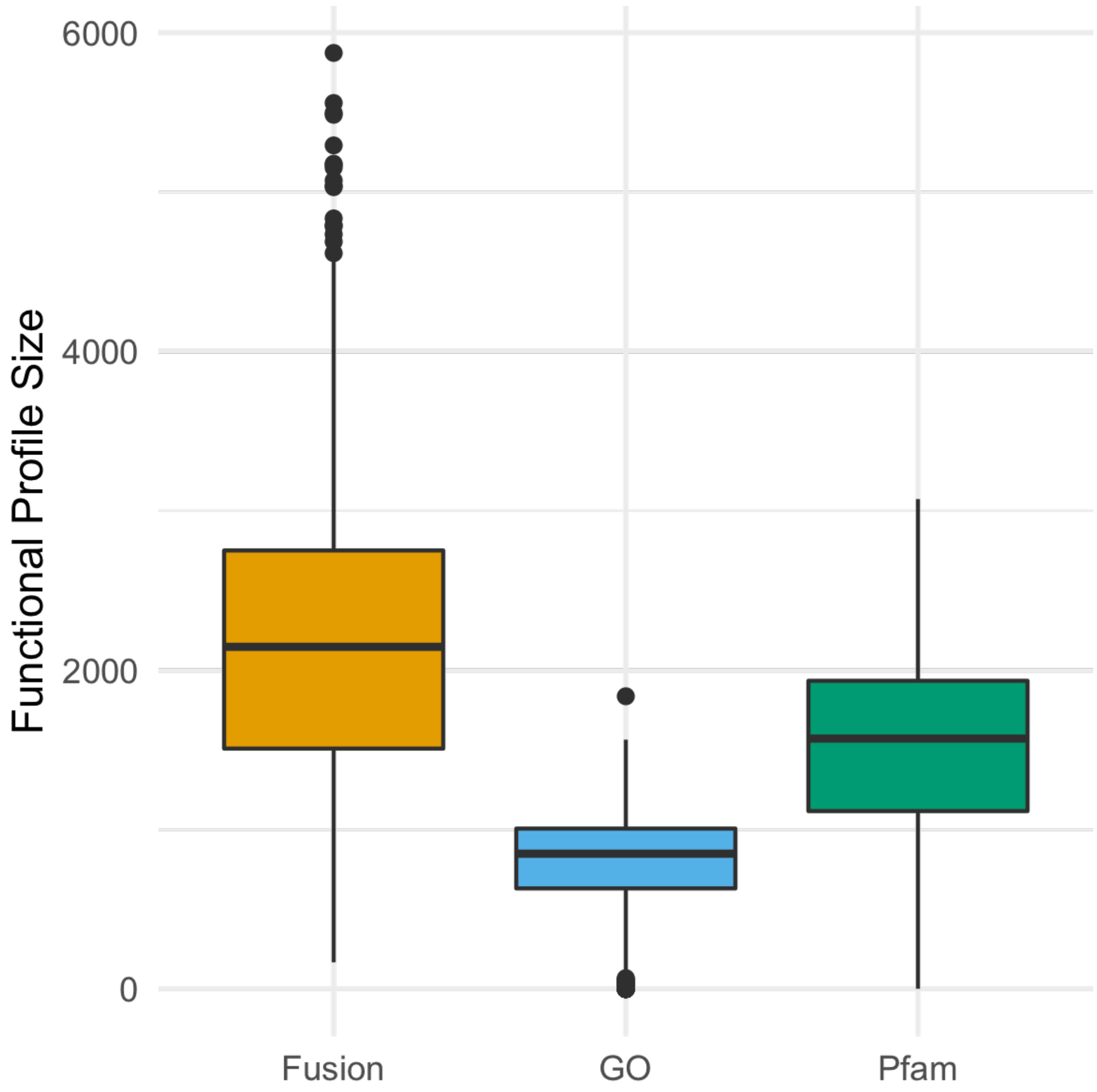


**Fig. S2. Average functional profiles based on Fusion are larger than Pfam and GO -based profiles**. Average, per organism Fusion (orange), Pfam-A (green) and GO term (blue) functional profiles are of size 2,133, 1,479, and 776 (median 2,147, 1,571, and 850), respectively. Minimal and maximal sizes are (168, 2, and 1) and (5,074, 3,178, 1,837), respectively.

**Fig. S3.**

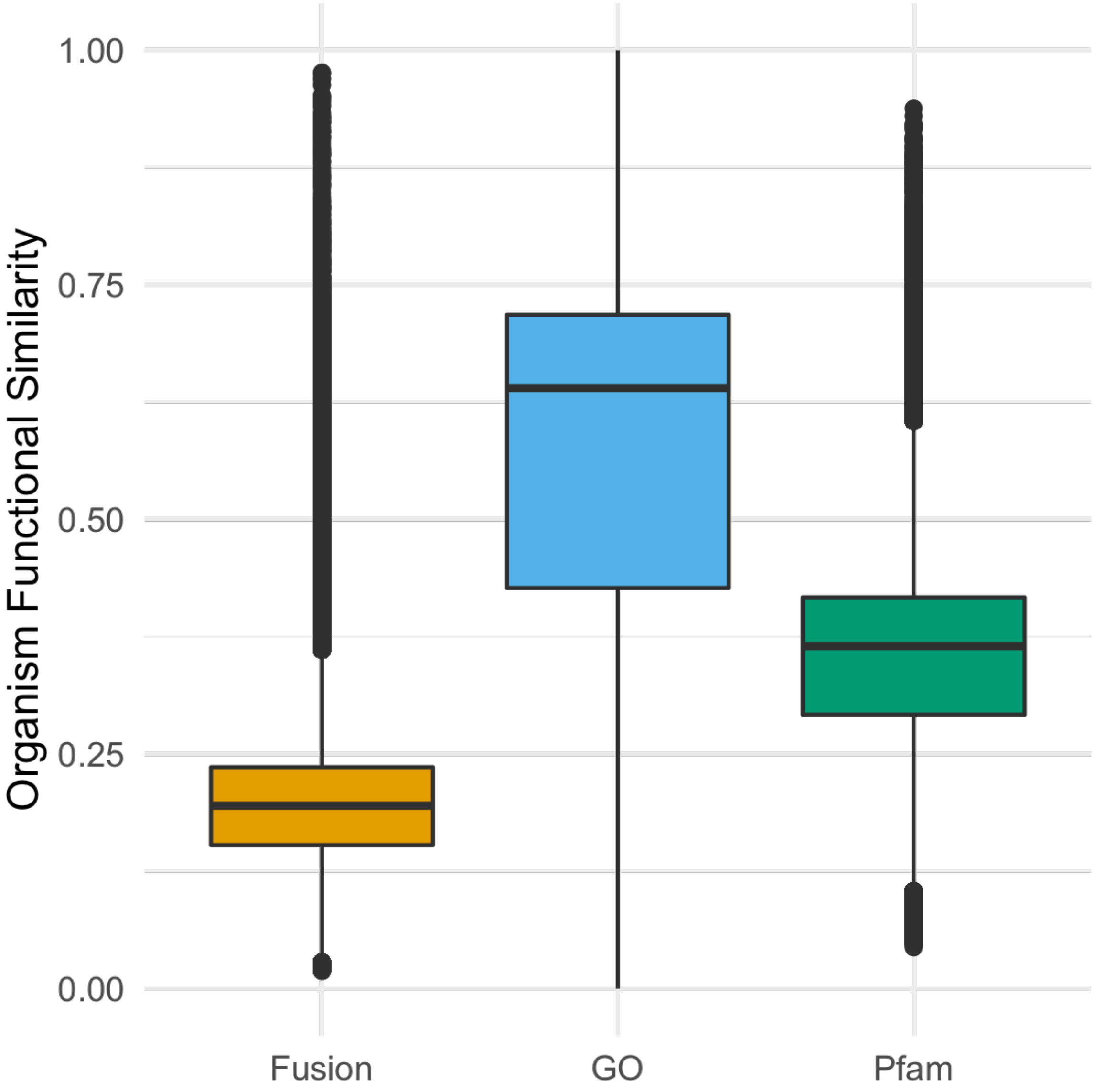


**Fig. S3. Larger functional profiles lead to lower organism similarities**. Organism pairwise similarities based on Fusion profiles (orange) ranged between 0.02 and 0.98 (median and mean similarity =0.20). For Pfam-A (green) and GO term (blue) based profile similarities min, max, median and mean similarity were (0.04, 0.94, 0.37, 0.35) and (0, 1, 0.64, 0.54) respectively. This is a directly inverse relationship with the organism functional profile sizes reported in Fig. S2.

**Fig. S4.**

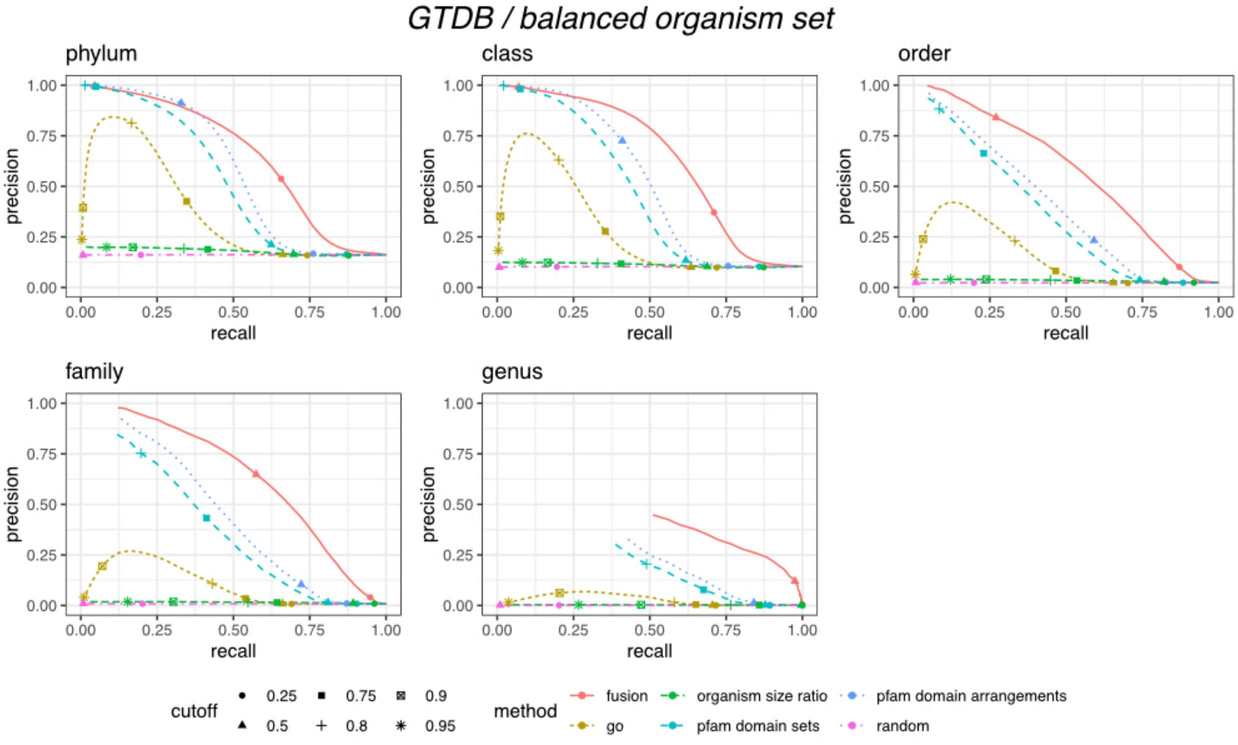


**Fig. S4. Fusion based organism similarity is best at defining pairwise organism taxonomic identity.** Shown in each panel is the precision (y-axis) at a given recall (x-axis) for correctly identifying if any two organisms share the same taxonomic rank. Precision/recall pairs for selected similarity cut-offs (0.25, 0.5, 0.75, 0.8, 0.9, 0.95) are indicated by different shapes. Displayed are only precision/recall values where predicted positive pairs (TP & FP) make up at least 0.1% of all possible pairs. Fusion (red solid line) performed significantly better than Pfam (dashed blue line) and GO (dashed olive) at identifying the organism pair taxonomic identity at all taxonomic levels (genus through phylum) defined by GTDB. Baseline performance is indicated by random (dashed purple) and functional profile size ratio (green) curves.

**Fig. S5.**

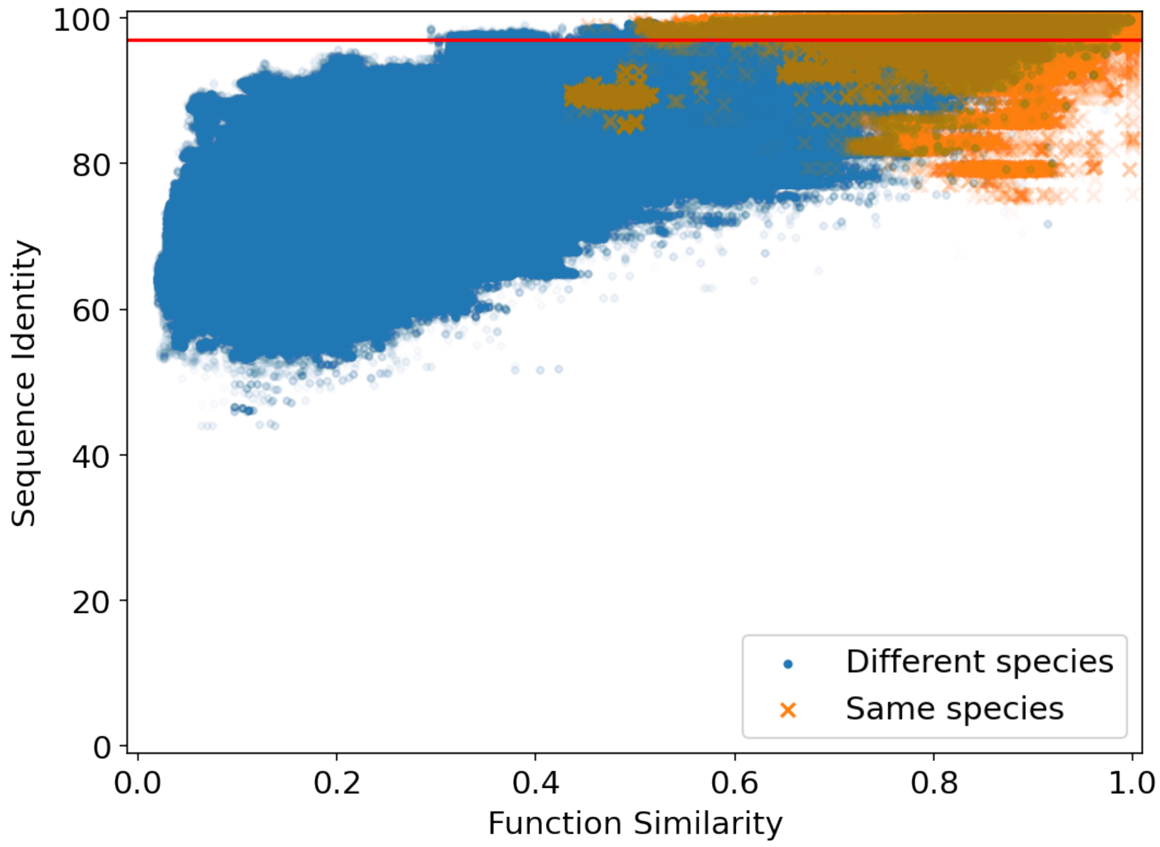


**Fig. S5. Fusion capable to reliably detect pairs of same taxonomic rank.** Shown are 16S rRNA sequence pairs of organisms of the same (orange crosses) vs different (blue dots) species. Organism pairwise 16S rRNA similarity, measured as sequence identity in a multiple sequence alignment (y-axis), is displayed in contrast to Fusion functional similarity (x-axis), measured as one minus the profile Jaccard distance. The red horizonal line at 97% represent the threshold below which 16S rRNA pairs are deemed to be originating from different species. Note that16S rRNA sequence identity and organism functional similarity appear to be correlated.

**Fig. S6.**

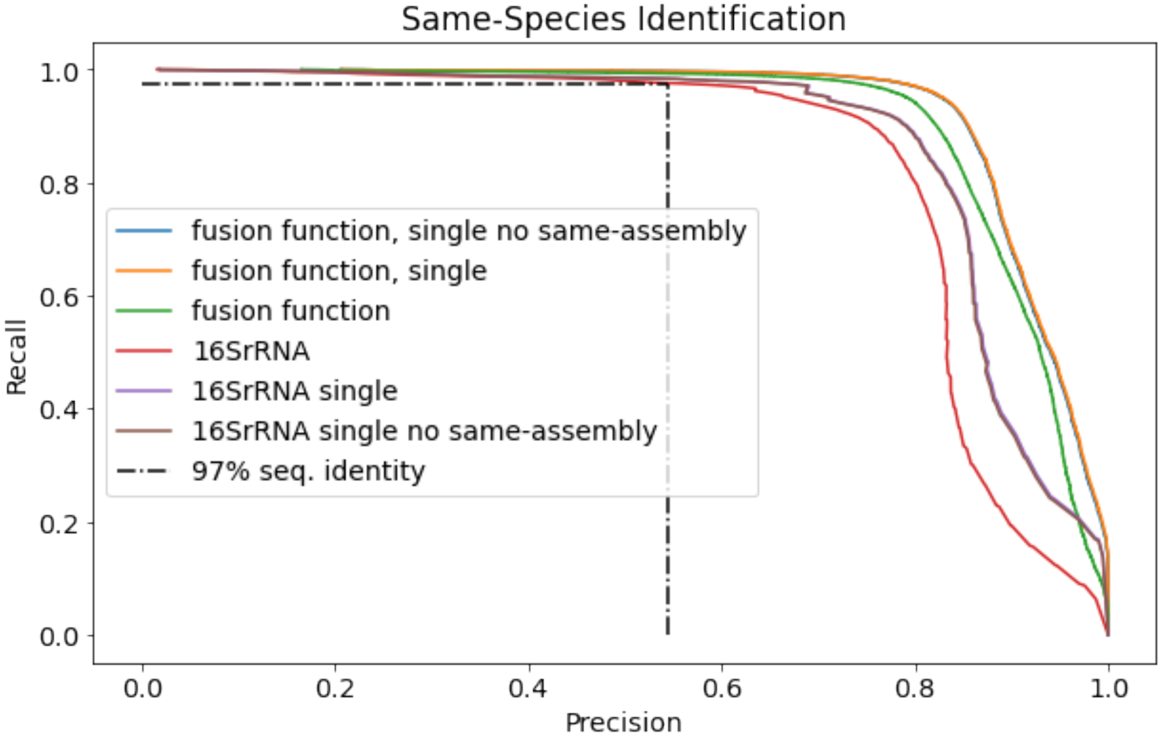


**Fig. S6. Fusion organism similarity outperforms 16S rRNA-based same-species identification.** Displayed are precision/recall values for correctly assigning 16S rRNA / organism pairs to the same species depending on their 16S rRNA and organism functional profile similarities. The 16S rRNA curve (red) represents all 16S rRNA pairs (potentially multiple per assembly). The corresponding green Fusion function curve is based on the same pairs, i.e. if multiple 16S rRNA pairings per assembly pair exist, the resulting organism functional profile is counted as many times in the precision recall calculation. Vice versa, "Fusion function, single" (orange) uses only one organism functional profile similarity per assembly pair; the corresponding "16S rRNA single" (purple) uses one, randomly chosen 16S pair per assembly pair. The "no same-assembly" curves (blue & brown) exclude assembly self-hits, i.e. pairs of organisms from the same assembly. Note that the orange and blue curves in the plot overlap.

**Fig. S7.**

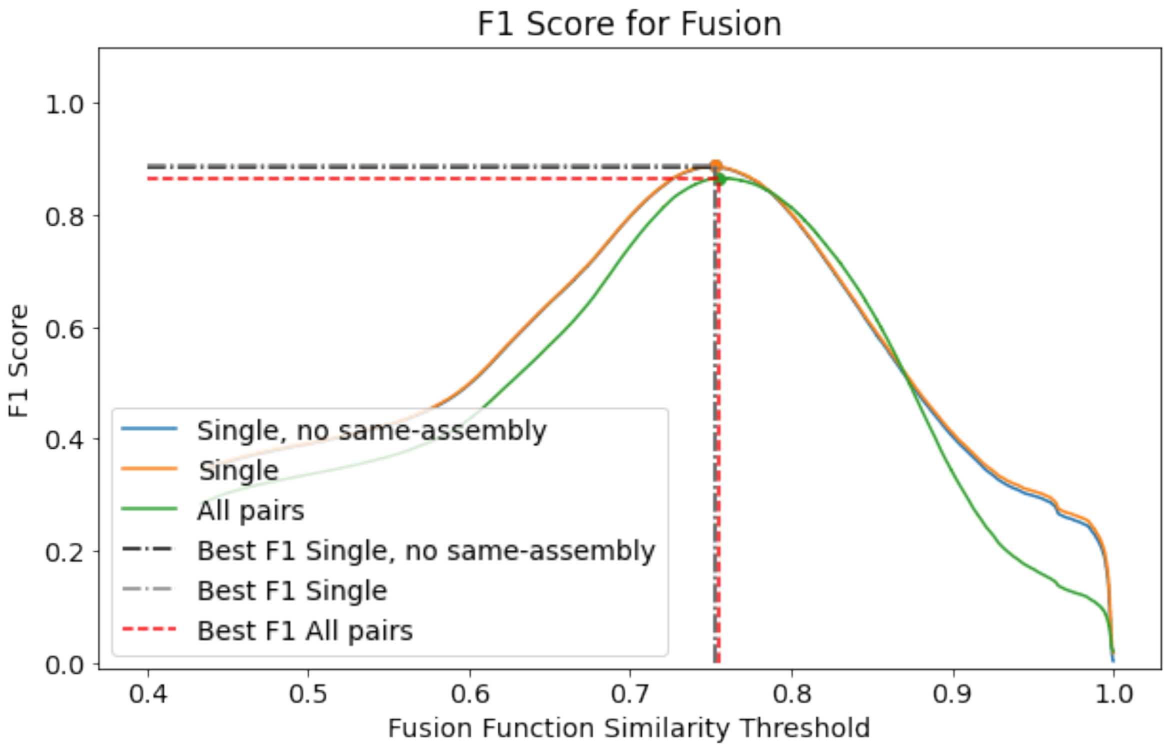


**Fig. S7. Selecting the optimal F1-score for same species detection using Fusion organism similarities (0.755).** Displayed is the F1-score (y-axis) plotted against the Fusion organism similarity threshold (x-axis) for same species assignment based on Fusion organism similarity. Single (orange) and "single, no same-assembly" (blue) curves consider only unique assembly pairs each (curves significantly overlap), whereas all pairs (green) is based on as many copies of organism similarities as 16S rRNA pairs exists for each given assembly pair.

**Fig. S8.**

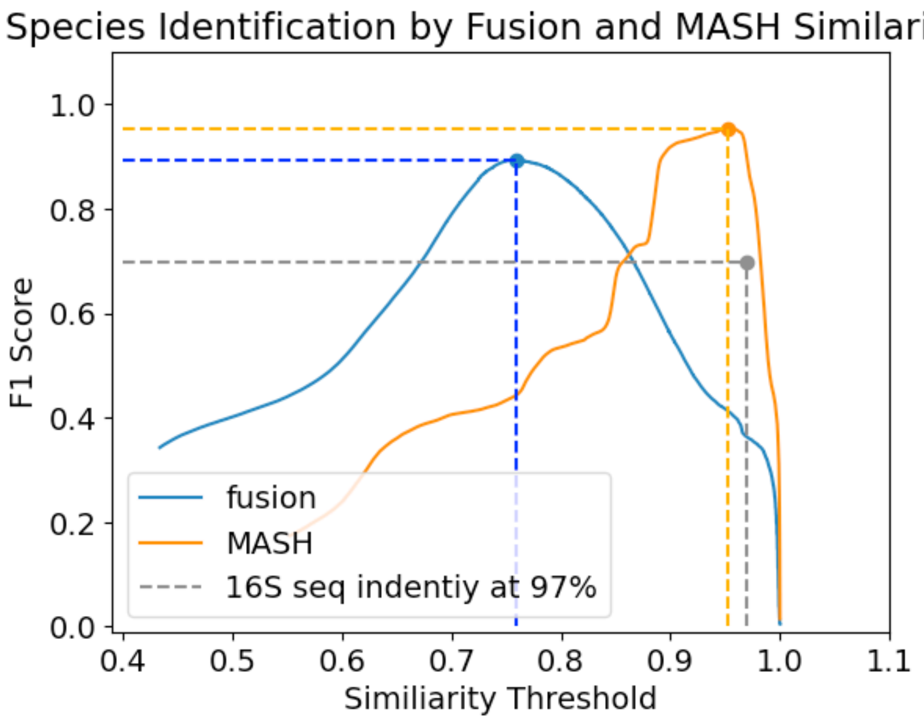


**Fig S8. Mash achieves slightly higher F1 Score than fusion.** Mash (orange) achieves an F1 score of ~0.95 at a similarity threshold of ~0.96, in comparison to fusion (blue, F1 score ~0.9 at 0.76 organism similarity). As reference at the default threshold for 16S rRNA (grey) of 0.97, the F1 Score is significantly lower (~0.7).

**Fig. S9.**

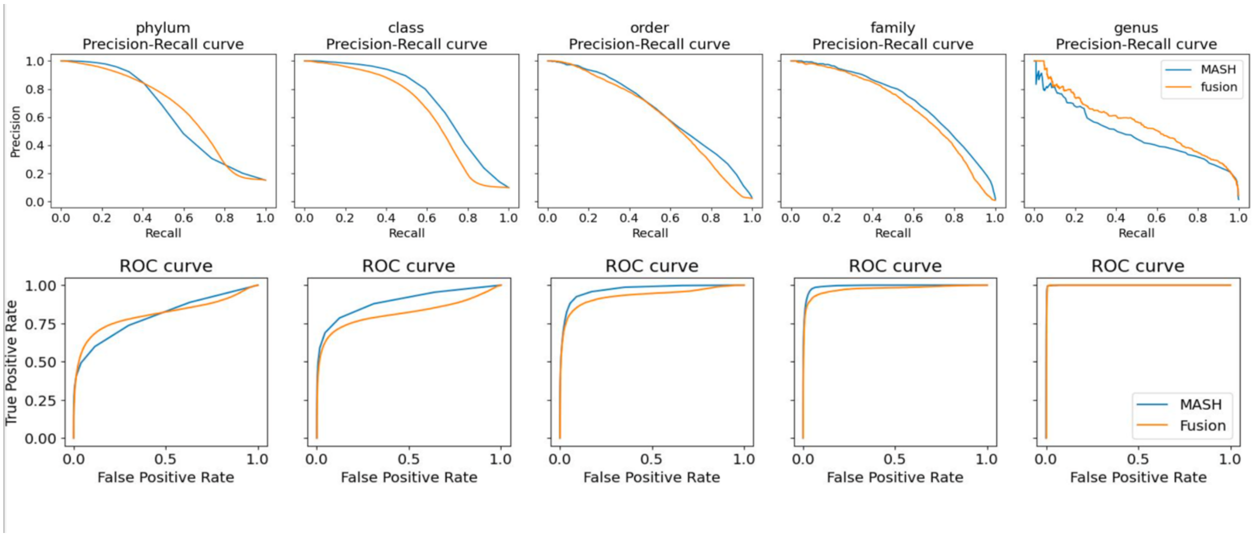


**Fig. S9. Mash performs similarly to fusion in identifying same taxonomic level for organism pairs.** Shown are Precision/Recall (top row) and ROC (bottom row) for identifying whether two organisms belong to the same taxonomic group (phylum through genus, columns). Performance is displayed for MASH based (blue) and fusion based (orange) identification. Organism pairs are chosen out of the balanced organism set.

**Fig. S10.**

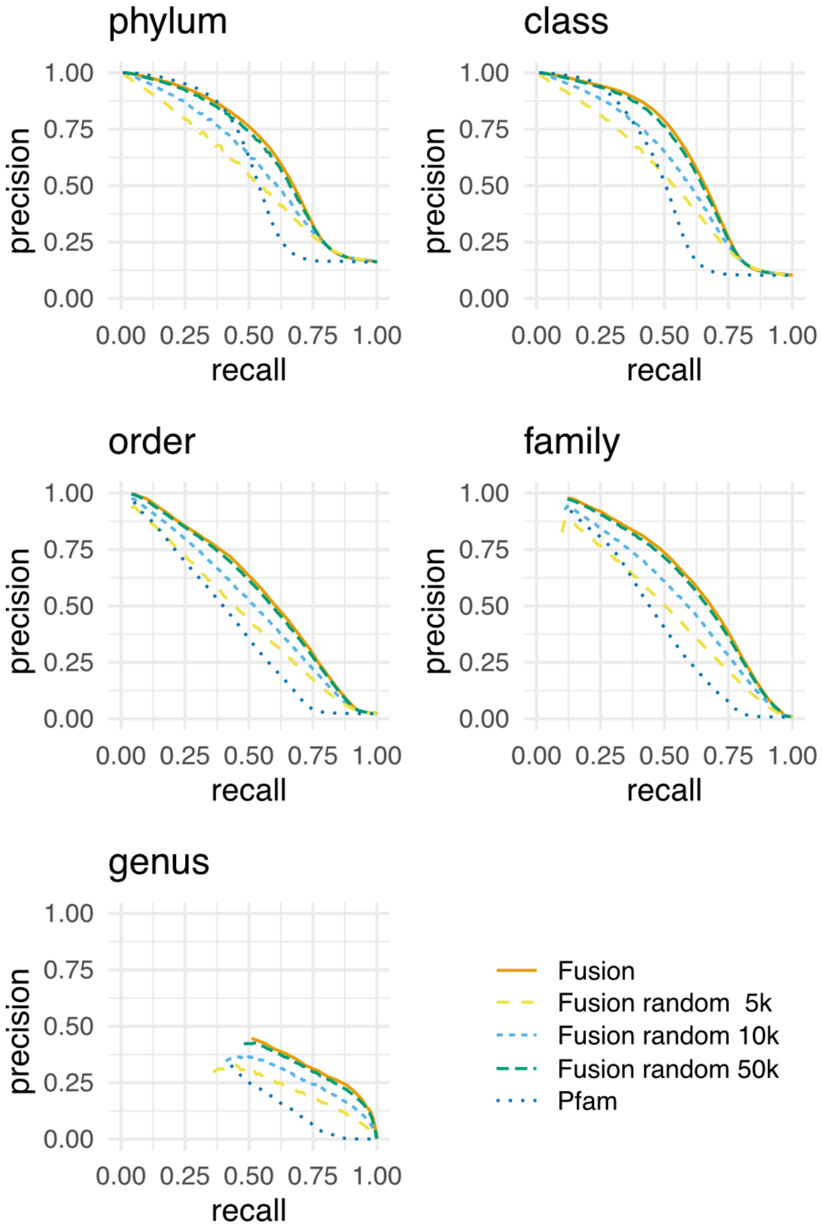


**Fig. S10. Randomly selected Fusion functions identify organism taxonomic relationships.** Each panel reflects the precision (y-axis) at a given recall (x-axis) for correctly identifying two organisms as sharing the same taxonomic rank (panel label). Line color indicates the functional samples. For example, using 5,000 Fusion functions (yellow) outperforms using all of Pfam domain arrangements (darkblue) for most cutoffs across all panels. Displayed are only precision/recall pairs where predicted positives pairs (TP+FP) make up at least 0.1% of all possible pairs**.**

**Fig. S11.**

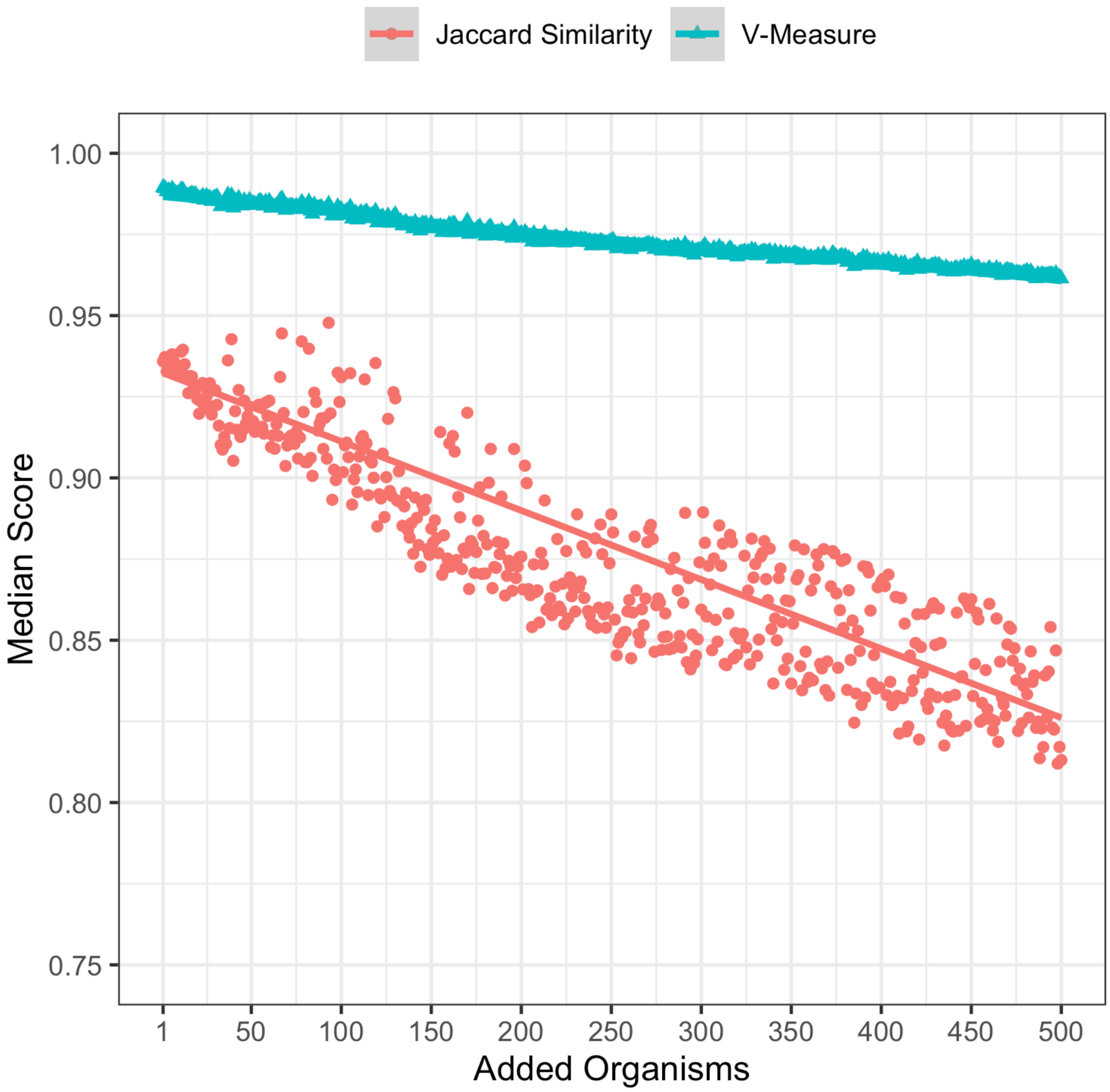


**Fig. S11. Fusion-informed taxa are stable with the addition of new organisms.** Additional organisms (1-500) are included in the generation of Fusion-informed taxa (additional to the organisms of the balanced organism set). With added organisms V-measure (turquoise) and Jaccard Similarity (red) marginally drop if compared to the reference of GTDB-taxon assignment.

**Fig. S12.**

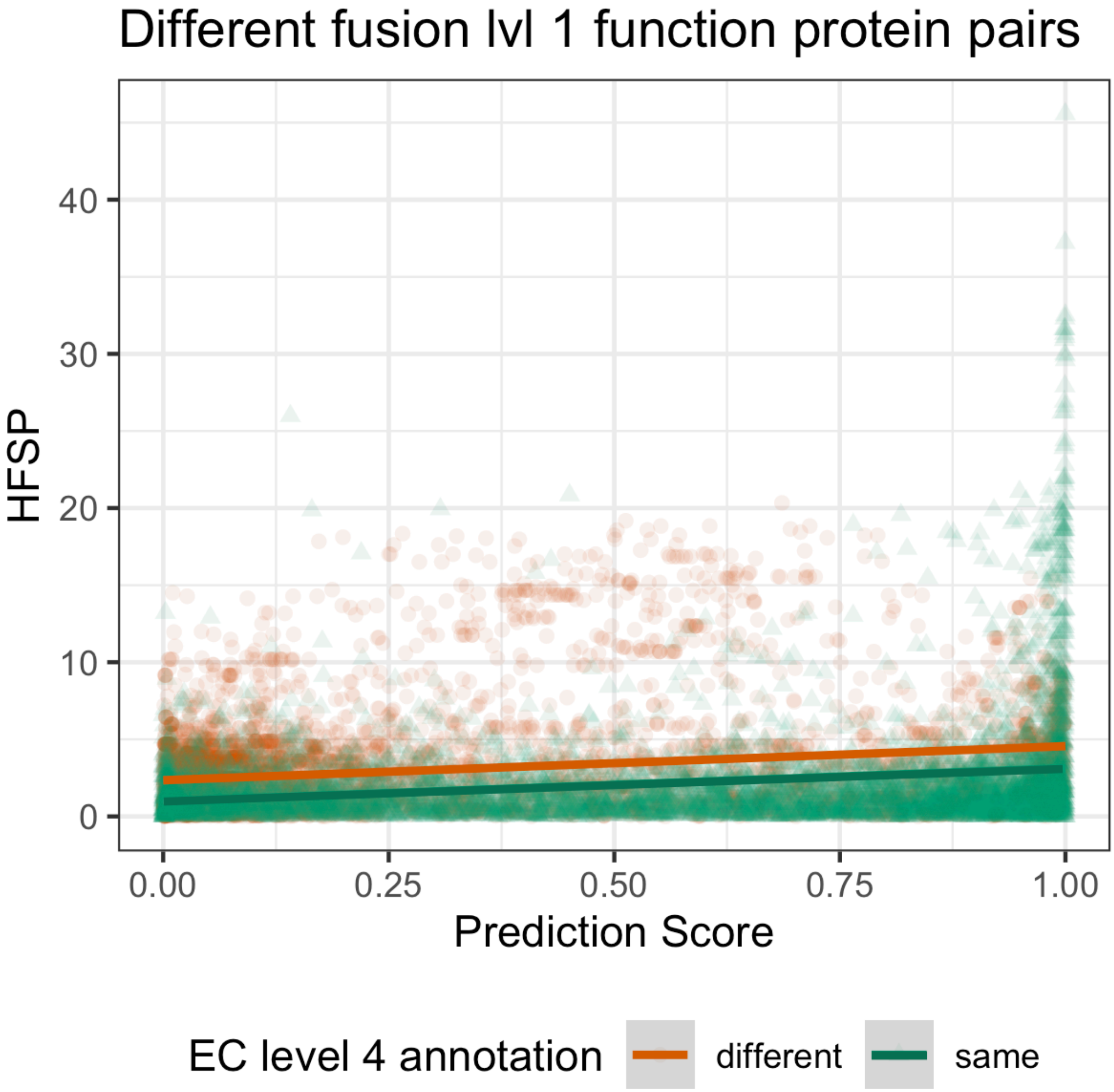


**Fig. S12. SNN-scores capture a different signal than sequence identity informed HFSP scores despite small correlation.** Displayed are SNN-scores (x-axis) and HFSP scores (y-axis) for protein pairs originating from different Functions. Each protein pair is coloured by whether they share the same or different EC annotation on the 4$^{th}$ level. Protein pairs mostly distribute evenly across the SNN-score spectrum independent from their HFSP score, with the exception of a slight increase in high HFSP / high SNN-scores for functionally very close protein pairs.

**Fig. S13.**

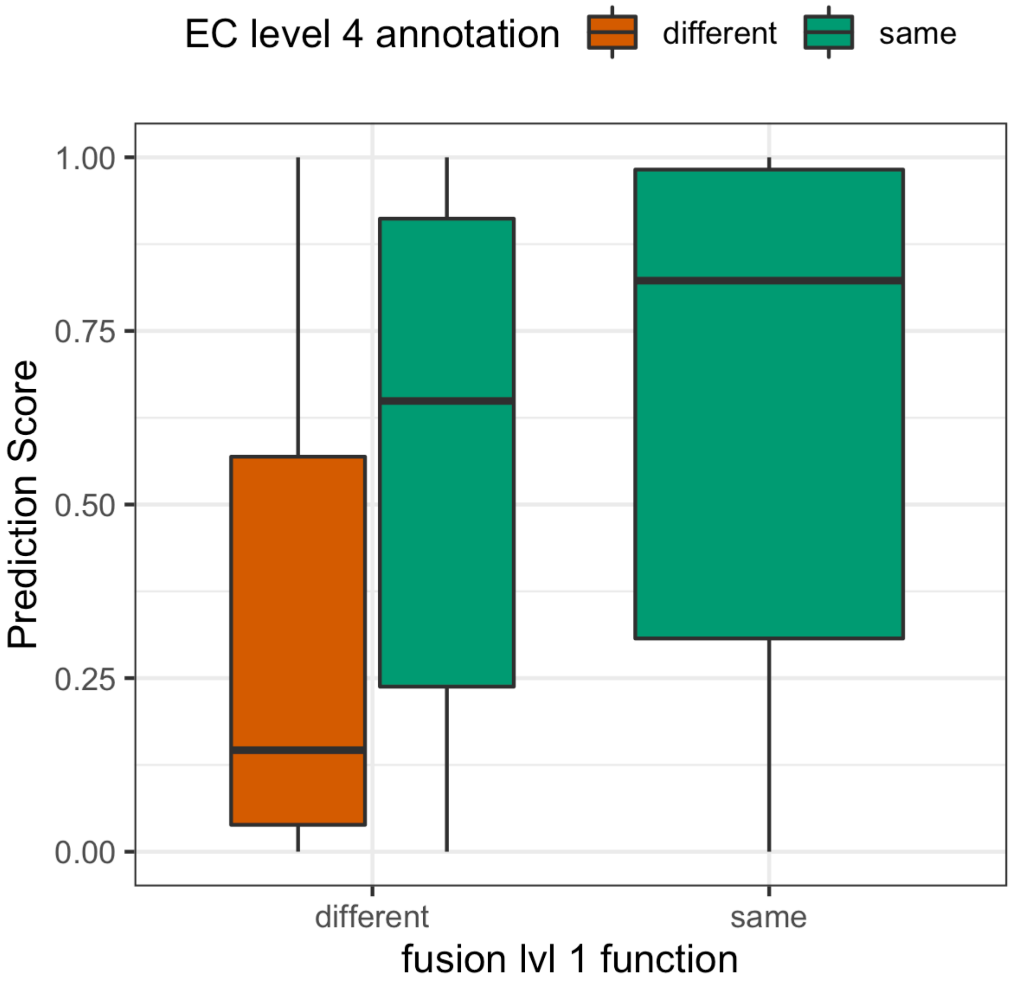


**Fig. S13. Protein pairs of different fusion function but with identical EC annotations are predicted with similar scores as proteins from the same fusion function.** Protein pairs with identical EC annotation (green) are predicted in similar score ranges independent from whether they are in the same fusion function or not. Protein pairs of differing fusion function and differing E.C. annotation (orange) are predominantly predicted with scores below 0.5 – contrary to protein pairs with the same E.C. annotation.

**Fig. S14.**

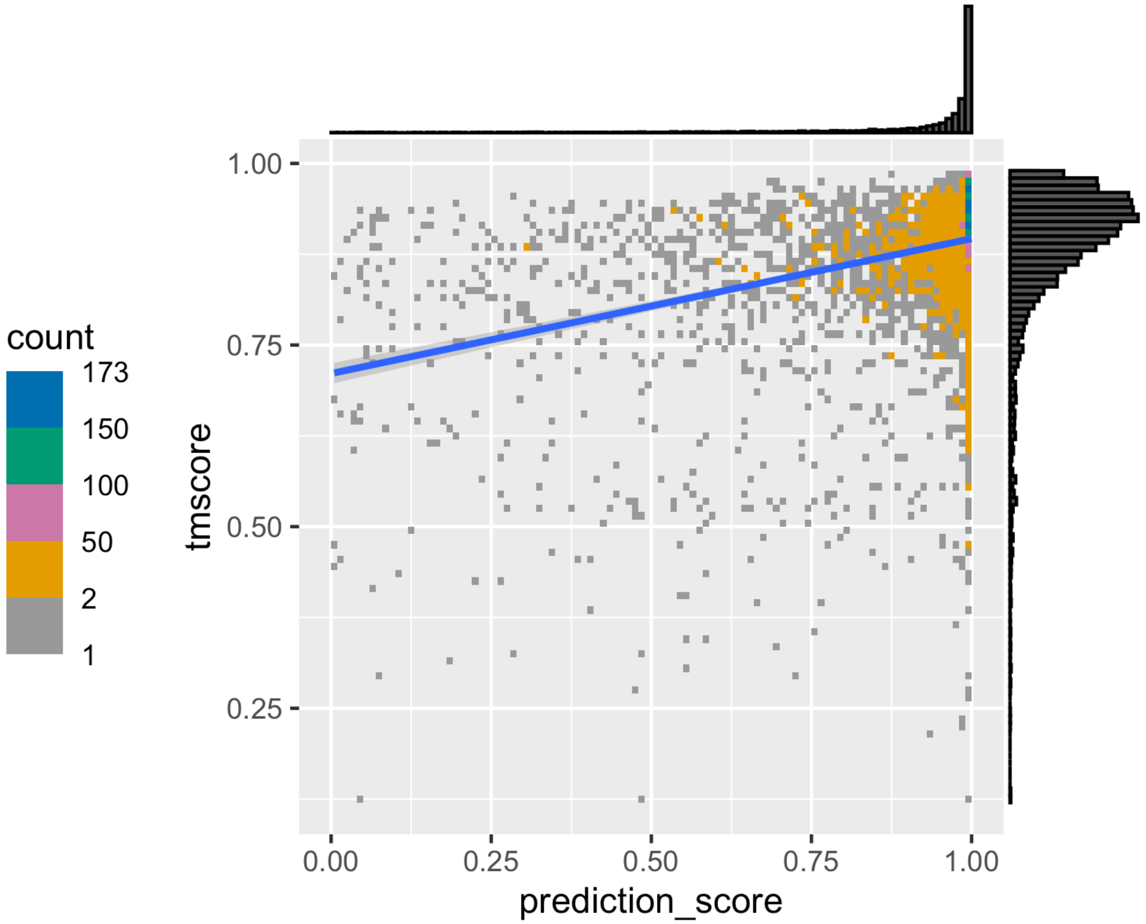


**Fig. S14. Protein pairs of functional similarity (HFSP≥0) have both high structural similarity as well as high SNN-score.** Displayed are only protein / PDB structure pairs that achieve an HFSP score ≥ 0. TM-score (y-axis) indicates structural similarity (with TM-score ≥ 0.7 generally regarded as same structure), SNN-score (x-axis) reports predicted functional similarity (high values indicate higher reliability for functional similarity). Nearly all protein / structure pairs for which an HFSP score was generated are predicted to be structurally / functionally by both TM-align as well as our SNN predictor.

**Fig. S15.**

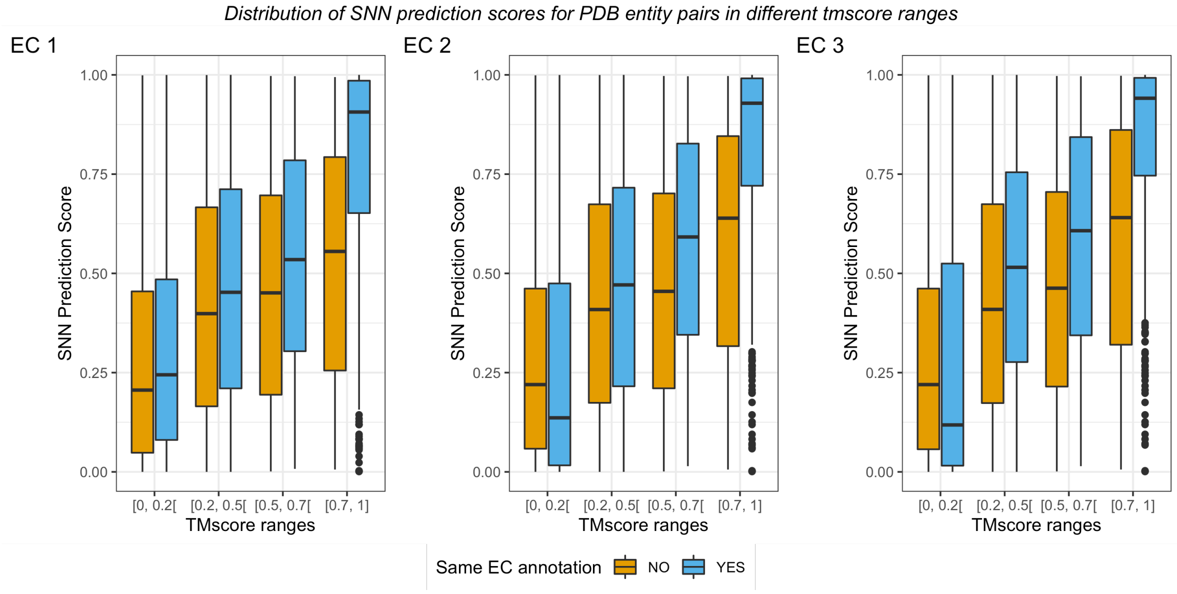


**Fig. S15. Protein / structure pairs with known identical EC annotation show higher SNN-scores within the same TM-score ranges than pairs with differing EC annotations.** Pairs of proteins / structures are grouped by EC annotation difference (blue = same, orange = different). For each EC level median SNN-scores increase with higher TM-score ranges. Within the same TM-score range protein pairs of same EC annotation tend to have higher SNN-scores than proteins of different EC annotation. This difference is most notable in the TM-score range of [0.7-1] which indicates high structural similarity.

**Table S1.**

| Taxonomic Group | Fusion clustering resolution | KS p-Value |
|---|---|---|
| Phylum | **0.68** | **0.89** |
| Class | **0.68** | **0.78** |
| Order | **0.5** | **0.68** |
| Family | **0.36** | **0.01** |

**Table S1. According to Kolmogorov-Smirnov test p-values, size distributions between GTDB taxa and corresponding Fusion-taxa are different phylum through order.** Fusion-taxa are generated by clustering the organism network given the listed Louvain Resolution thresholds. KS-test p-values $< 0.05$ indicate that distributions of taxa sizes are similar.

**Table S2.**

| | HFSP ≥ 0: NO | HFSP ≥ 0: YES |
|---|---|---|
| 0.7 ≤ TM-score ≤ 1 | 13,874<br>0.2% | 3,988<br>0.0% |
| 0.5 ≤ TM-score < 0.7 | 39,452<br>0.5% | 90<br>0.0% |
| 0.2 ≤ TM-score < 0.5 | 363,486<br>4.2% | 16<br>0.0% |
| 0 ≤ TM-score < 0.2 | 29,531<br>0.3% | 0<br>0.0% |
| No TM-align | 8,150,403<br>94.8% | 38<br>0.0% |

**Table S2.** Counts of structure pairs in TM-score range (rows) separated by whether they are functionally similar as by HFSP (columns). Percentage values are calculated across the whole table; shading is performed row-wise, i.e. darker shades of blue indicate larger values within one row.

**Table S3.**

| | HFSP ≥ 0: NO | HFSP ≥ 0: YES |
|---|---|---|
| 0.98 ≤ SNN-score ≤ 1 | 155,117<br>1.8% | 2,119<br>0.0% |
| 0.5 ≤ SNN-score < 0.98 | 3,538,115<br>41.1% | 1,757<br>0.0% |
| 0 ≤ SNN-score < 0.5 | 4,903,514<br>57.0% | 256<br>0.0% |

**Table S3.** Counts of protein pairs in SNN-score range (rows) separated by whether they are functionally similar as by HFSP (columns). Percentage values are calculated across the whole table; shading is performed row-wise, i.e. darker shades of blue indicate larger values within one row.

**Table S4.**

| | 0 ≤ SNN-score < 0.5 | 0.5 ≤ SNN-score < 0.98 | 0.98 ≤ SNN-score ≤ 1 |
|---|---|---|---|
| 0.7 ≤ TM-score ≤ 1 | 4,112<br>0.0% | 10,167<br>0.1% | 3,583<br>0.0% |
| 0.5 ≤ TM-score < 0.7 | 18,455<br>0.2% | 19,456<br>0.2% | 1,631<br>0.0% |
| 0.2 ≤ TM-score < 0.5 | 201,370<br>2.3% | 155,020<br>1.8% | 7,112<br>0.1% |
| 0 ≤ TM-score < 0.2 | 21,668<br>0.3% | 7,782<br>0.1% | 81<br>0.0% |
| No TM-align | 4,658,165<br>54.2% | 3,347,447<br>38.9% | 144,829<br>1.7% |

**Table S4.** Counts of structure pairs in TM-score range (rows) separated by whether they are functionally similar as by SNN-score (columns). Percentage values are calculated across the whole table; shading is performed row-wise, i.e. darker shades of blue indicate larger values within one row.


1. D. A. Benson *et al.*, GenBank. *Nucleic Acids Res* **41**, D36-42 (2013).
2. E. W. Sayers *et al.*, GenBank. *Nucleic Acids Res* **47**, D94-d99 (2019).
3. G. A. Blackwell *et al.*, Exploring bacterial diversity via a curated and searchable snapshot of archived DNA sequences. *PLOS Biology* **19**, e3001421 (2021).
